# Integrating brain structure and function for the neurobiology and genetics of language

**DOI:** 10.64898/2025.12.17.694832

**Authors:** Jitse S. Amelink, Sourena Soheili-Nezhad, Gökberk Alagöz, Alberto Llera, Dick Schijven, Meng-Yun Wang, Koen V. Haak, Simon E. Fisher, Christian F. Beckmann, Clyde Francks

## Abstract

Brain structure and function have largely been studied separately in relation to the neurobiology and genetics of language. Here we used linked independent component analysis to integrate language network functional connectivity with brain volumetric and white matter structure in 32,677 UK Biobank participants, followed by analysis of behavioural, neurobiological and genetic correlates of the derived multimodal structure-function imaging components. Stronger functional connectivity between brain language areas was associated with increased volume of parts of the cerebellum and motor cortex, together with smaller ventricles and sensory parietal and occipital areas. The brain structure-function language components mediated an association between vocabulary level and polygenic scores for reading ability. We report 18 genomic loci associated with brain structure-function language components. Single-nucleotide polymorphism (SNP)-based heritability estimates for these components were 23-30%, and there was significant enrichment of heritability in primate-conserved genomic loci and fetal brain human-gained enhancer elements. This study revealed that structural correlates of functional language network connectivity extend well beyond previously defined language areas of the brain, and highlights the value of multimodal brain phenotyping for human neurogenetic discovery.

## Introduction

Key areas of the brain involved in language, in the left-hemisphere inferior frontal lobe and superior temporal lobes (classically referred to as Broca’s and Wernicke’s regions respectively), have been identified through lesion mapping studies^1^ and functional imaging^2–4^. Although notable exceptions exist^5–8^, most neurobiological studies of language have focused either on brain structure or function alone. This distinction is partly artificial, as structural and functional organization are closely intertwined. Brain structure necessarily constrains and sub-serves function^9,10^, while function also affects structure through plasticity-dependent processes in development, learning and aging^11–16^. Accordingly, genetic correlations between diverse brain structural and functional measures have been reported to range from roughly 0.2 to 0.6^17^, which indicates partial overlap in their genetic architectures.

Brain imaging studies have revealed substantial individual differences in structural and functional connectivity between key regions of the cerebral cortex involved in language^18–21^. Accordingly, recent large-scale genome-wide association studies (GWAS) of structural connectivity^22^ or functional connectivity^23,24^ between key language regions have been performed, in which individual differences in language network organization were linked to common genetic polymorphisms in the general population. These studies revealed clues to the molecular basis of brain organization in support of language. Indeed, 6 of 14 genomic loci associated with functional connectivity between core language areas^23^ were also found to affect structural connectivity^22^ which underscores the biological coupling of structure and function. However, thus far no genetic study of the brain’s infrastructure for language has integrated both functional and structural imaging modalities to derive multimodal phenotypes.

While there is evidence that language is processed by a distinct modular brain network that can be clearly discriminated from the rest of an individual’s brain^3^, other work has suggested strong integration of language with, for example, memory^2,25^ or motor organization^26^. In the present study we hypothesized that individual differences in the functional organization of inferior frontal and middle temporal regions may covary to an extent with functional and structural variation more broadly across the brain. We used linked independent component analysis (linked ICA^10,27^) to identify data-driven, multimodal imaging components that capture individual differences shared across both structure and function in 32,677 adults in the UK Biobank, to reveal undiscovered aspects of the neurobiology and genetics of language.

To first contextualize the multimodal structure-function language components, we compared their corresponding brain maps in each contributing modality against maps for cognitive functions and neurotransmitter densities maps (using NiMARE^28^, Neurosynth^29^ and Neuromaps^30^). To more specifically assess relevance to language-related cognition, we extracted individual participant contributions to the multimodal components and tested for relations with vocabulary levels (see Methods for a description) and polygenic dispositions to language-related traits and neurodevelopmental conditions. We then used the individual participant contributions as imaging-derived phenotypes for GWAS and SNP-based heritability analysis. Post-GWAS analysis consisted of gene mapping, gene-set analysis for brain development and cell-types, SNP-based enrichment analysis for cell-types, and transcriptome-wide association studies (TWAS) for dissecting the genetic and neurobiological architecture. Finally, as language is a uniquely human form of communication in its degree of complexity and functionality within the animal kingdom, we examined genetic contributions to the multimodal brain imaging components with respect to various annotations derived from evolutionary genomic analysis, including conservation across primate species, and human-gained fetal brain enhancers. See Figure 1 for an overview of the study.

**Figure 1:**
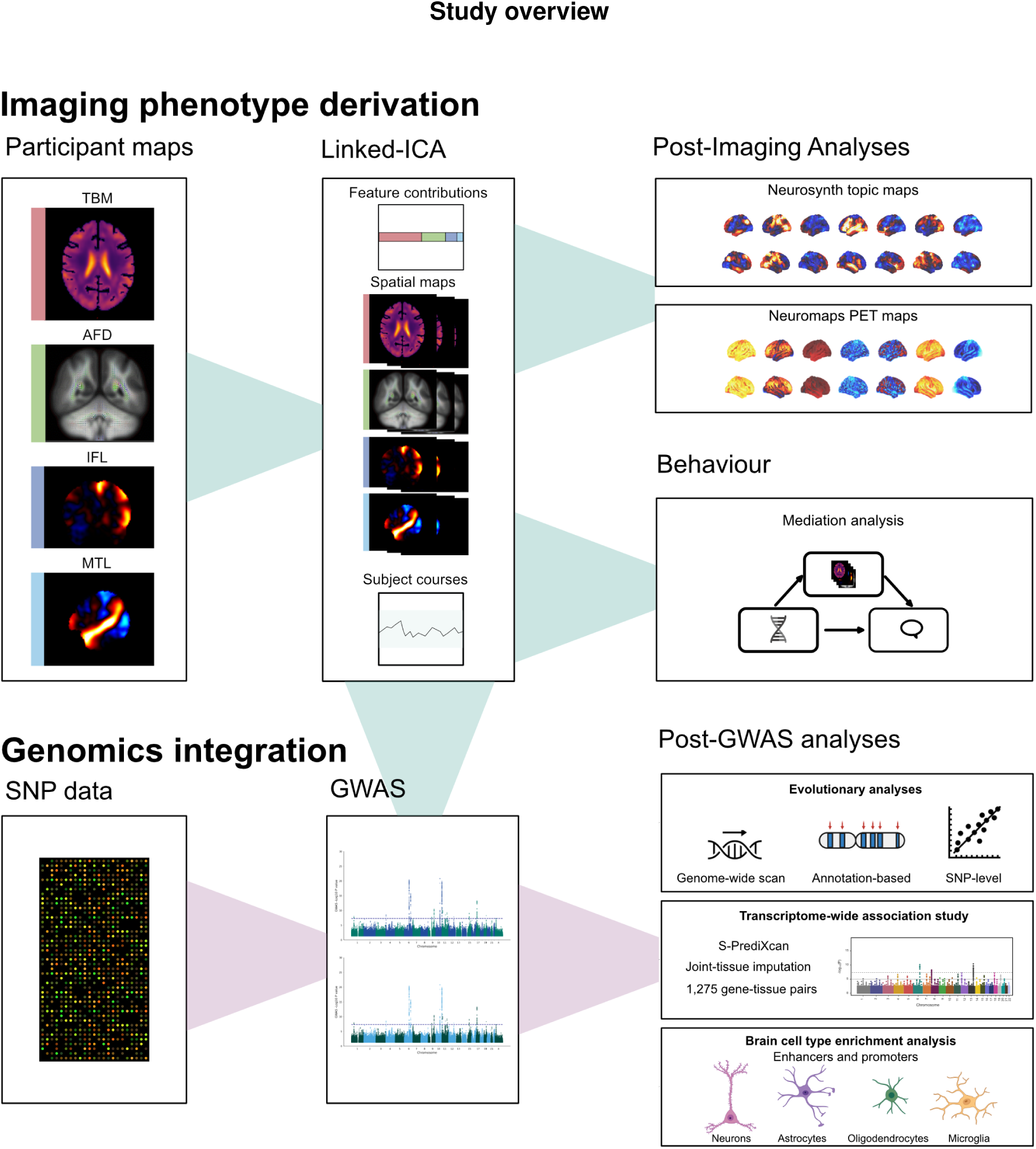
Overview of the study. We derived unimodal brain maps for 32,677 participants in UK Biobank for grey matter (TBM), white matter (AFD) and resting state functional connectivity (IFL and STL) and entered these into linked-ICA, then selected multimodal imaging components with a functional contribution. We subsequently decoded the corresponding brain maps to these components using cognitive and molecular annotations, and investigated associations between vocabulary levels and polygenic scores for various language-related traits and conditions. Multimodal components were also used as phenotypes for GWAS and heritability analysis, followed by gene mapping and gene set-analysis, transcriptome-wide association, cell-type analysis, and partitioned heritability analysis with respect to genomic annotations for evolutionary conservation and selection.

## Results

### Broad and narrow structural correlates of core brain language areas extend beyond canonical language regions

For 32,677 participants from the UK Biobank (adult general population dataset) we applied a Bayesian data-fusion algorithm called linked-ICA^27^ to 4 MRI-derived brain maps from three different imaging modalities: whole-brain volumetrics (derived using tensor-based morphometry, TBM^31^), whole-brain white matter apparent fibre density (AFD, derived using fixel-based analysis,^31,32^), and whole-brain resting state functional connectivity maps for the inferior frontal lobe (IC 28, derived using FSL by UK Biobank^33,34^) and superior temporal lobe (IC 9, derived using FSL by UK Biobank^33,34^). We derived 5 component (5c) and 10 component (10c) solutions, which correspond to lower and upper optima based on scree plots from principal component analysis of each MRI modality separately (Supplementary Figures 1-3). The 5c and 10c decompositions produced one component each with major contributions from functional connectivity for core language network regions, but also capturing structural variation (> 0.4 on multimodality index, minimum resting state contribution > 0.2 on unimodal resting state maps, and individual participant dominance < 0.01).

In particular, an extensive set of regions in the TBM structural modality is captured by the 5c multimodal comonent, which we henceforth refer to as the “broad language component”, whereas a more narrowly defined set of regions in the TBM structural map is captured by the 10c multimodal component, and we refer to this here as the “narrow language component”. As expected, for both the broad and narrow language components, strong resting state contributions from the inferior frontal lobe (IFL) (broad: 33.5%, narrow: 63.6%) and superior temporal lobe (STL) (broad: 18.3%, narrow: 34.0%) are present (Figure 2). Both components display left-lateralized patterns in IFL and to a lesser extent in STL (Figure 2), consistent with left-hemisphere language dominance^35^ being the typical organization in the population. The frontal brain map also extends to the supplementary motor area, whereas the superior temporal brain map extends throughout the temporal lobe.

**Figure 2:**
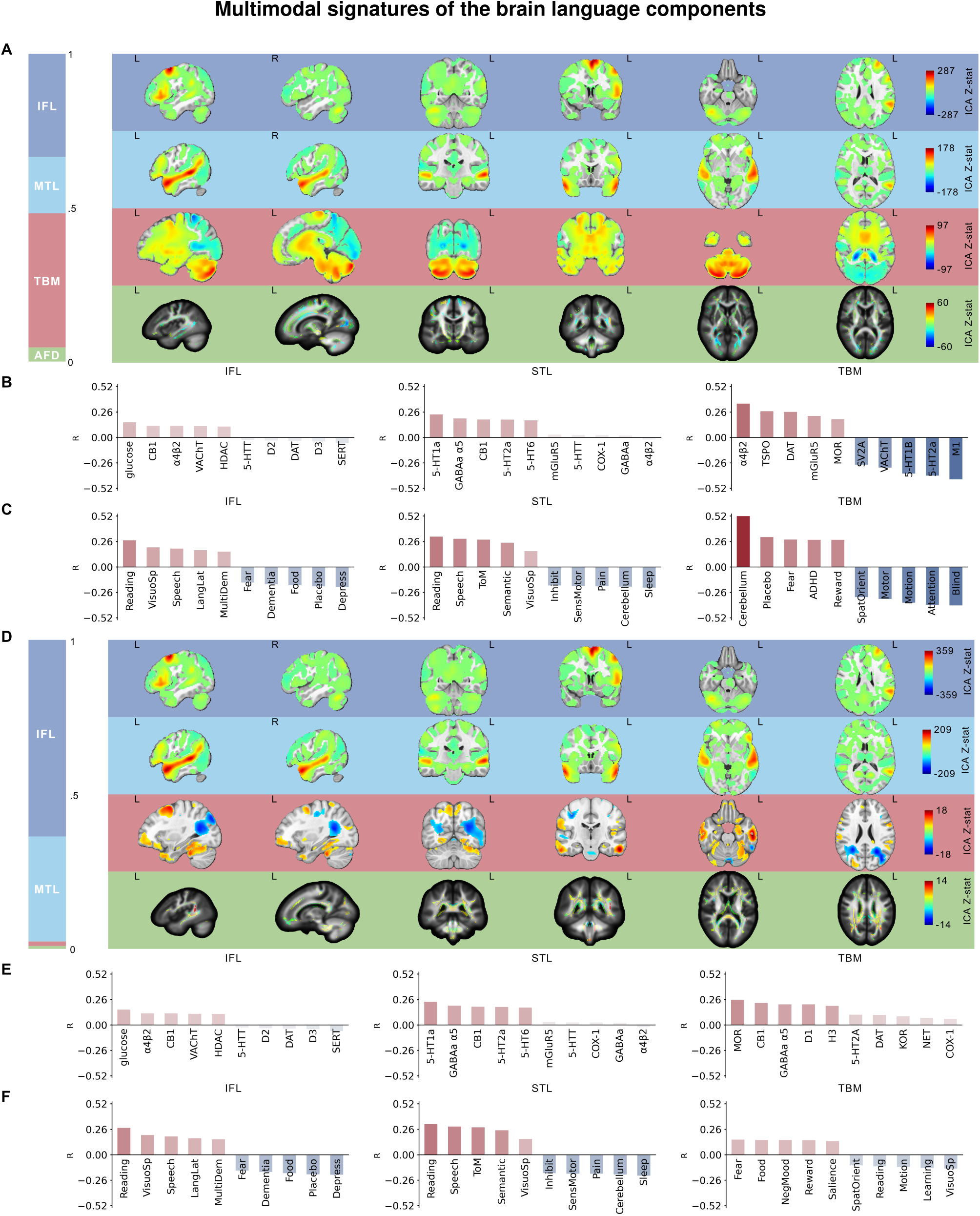
The broad **(A-C)** and narrow **(D-F)** components that capture multimodal individual differences linked to language network connectivity. **A+D.** On the left proportions of modality contributions are shown, on the right brain maps from top to bottom: resting state inferior frontal lobe (IFL), resting state superior temporal lobe (STL), tensor-based morphometry (TBM) and apparent fibre density (AFD). **B+E** Decoding of respective TBM brain maps with respect to meta-analyzed functional term data (Neurosynth). **C+F.** Cortical neurotransmitter densities for TBM brain maps (using NiMARE and neuromaps: see Methods).

The broad structural correlates of the language network consist of a broad positive-negative mode that runs anterior-to-posterior in the brain, with a large contribution from the TBM modality (43.5%) and a smaller contribution from AFD (4.6%), resulting in a high degree of multimodality (MMI = 0.75). Increased resting state connectivity in the IFL and the STL are associated with increased volume of cerebellar and prefrontal areas (Figure 2, Supplementary Figure 4), but reduced volume in sensory areas, ventricles and the optic radiation (both in TBM and AFD, Figure 2). The narrow structural correlates display less multimodality (MMI = 0.48) with smaller contributions from both TBM and AFD modalities (both around 1%). The resulting spatial patterns are more localized to the orbitofrontal cortex and parts of the temporal lobe (Figure 2).

### Contextualizing the multimodal component maps

To understand the cognitive and neurotransmitter architectures, we decoded our brain maps against Neurosynth topics (50 topics, v7)^29^ and neurotransmitter PET maps^30^ using NiMARE^28^. As expected, both the broad and narrow IFL and STL functional brain maps are prominently associated with language-related topics such as ‘Language and reading’ and ‘Auditory/speech’ (Figure 2, Supplementary Table 1). Apart from language-related annotations, the resting state IFL brain maps correlate with visuospatial orientation, whereas the resting state STL brain maps correlate with the theory of mind network. The extended structural correlates show a distinct pattern from either of the resting state maps (Figure 2, Supplementary Table 1). The strongest positive associations are observed with cerebellar function, and fear and reward processing, while the strongest negative associations with blindness, attention and various sensorimotor topics (motion, spatial orientation and motor).

Molecular profiles associated with the broad and narrow resting state IFL maps include cannabinoid receptor 1 (CB1), the vesicular acetylcholine transporter (VAChT), *α*4*β*2 nicotinic acetylcholine receptor, and glucose metabolism. The STL broad and narrow resting state maps on the other hand have their strongest similarity with three serotonergic maps (5-HT1a, 5-HT2a, 5-HT6) as well as cannabinoid (CB1) and GABAergic (GABAa *α*5) maps (Figure 2, Supp Table 2). 5-HT1a was previously found to be more strongly expressed in language areas compared to primary sensory areas^36,37^.

For the broad language component, structural associations with molecular maps display a notable opposite pattern to those of resting state maps (Figure 2, Supplementary Table 2), showing negative correlations with cholinergic M1 receptors, several serotonergic maps (most notably 5-HT1b and 5-HT2a) and kainate receptor maps (mGluR5). The cholinergic system showed associations in opposing directions in relation to broad language component structure: a strong positive association with *α*4*β*2 receptors and strong negative association with VAChT. For the narrow language component, structural associations with molecular maps showed heterogeneous patterns, with the strongest associations with mu-opioid receptors (MOR) and cannabinoid receptors (CB1) (Figure 2, Supplementary Table 2).

### Brain language components mediate an association between vocabulary level and polygenic scores for reading ability

We used individual participant contributions to the multimodal components as phenotypes for subsequent analyses. To understand whether the multimodal brain language components relate to language-related performance at the behavioural level, we tested their associations with vocabulary levels in 21,195 UK Biobank participants having these data available. Note that vocabulary level, as assessed with a picture test of receptive vocabulary (Methods), is currently the only cognitive test with close relevance for language available from the UK Biobank in more than 5,000 individuals. Both the broad (*β* = 0.039, *P* = 4.4 * 10^−8^) and narrow (*β* = 0.029, *P* = 3.9 * 10^−5^) language components showed subtle but significant associations with vocabulary levels.

To understand whether the broad and narrow language components in the brain might associate with genetic dispositions to further language-related traits, in the same set of 21,195 UK Biobank participants we generated polygenic scores (PGS) for reading ability^38^, left-handedness^39^ (which associates weakly with hemispheric language dominance^35^), educational attainment^40,41^, the reading disorder dyslexia (a condition involving difficulties with reading and spelling)^31,42^, autism^43^ (in which language ability is often reduced^44^ and reduced language lateralization has been reported^45^), ADHD^46^ (attention deficit/hyperactivity disorder, which often co-occurs with dyslexia^47^), and schizophrenia^48^ (in which language can be disordered and auditory hallucinations can occur^49^). Only the PGS for reading ability is associated with the multimodal brain language components after multiple testing correction (Figure 3, Supplementary Table 3). We also found that all of the polygenic scores, except the polygenic score for left-handedness, are associated with vocabulary level (Figure 3, Supplementary Table 3). Given this overall pattern of results, we conducted a mediation analysis with vocabulary level as the dependent variable, polygenic score for reading performance as predictor, and brain structure-function language components as potential mediators in the 21,195 participants. Both the broad and narrow brain language components mediate small but significant parts of the association between reading-ability PGS and vocabulary level (Figure 3, Supplementary Table 3).

**Figure 3:**
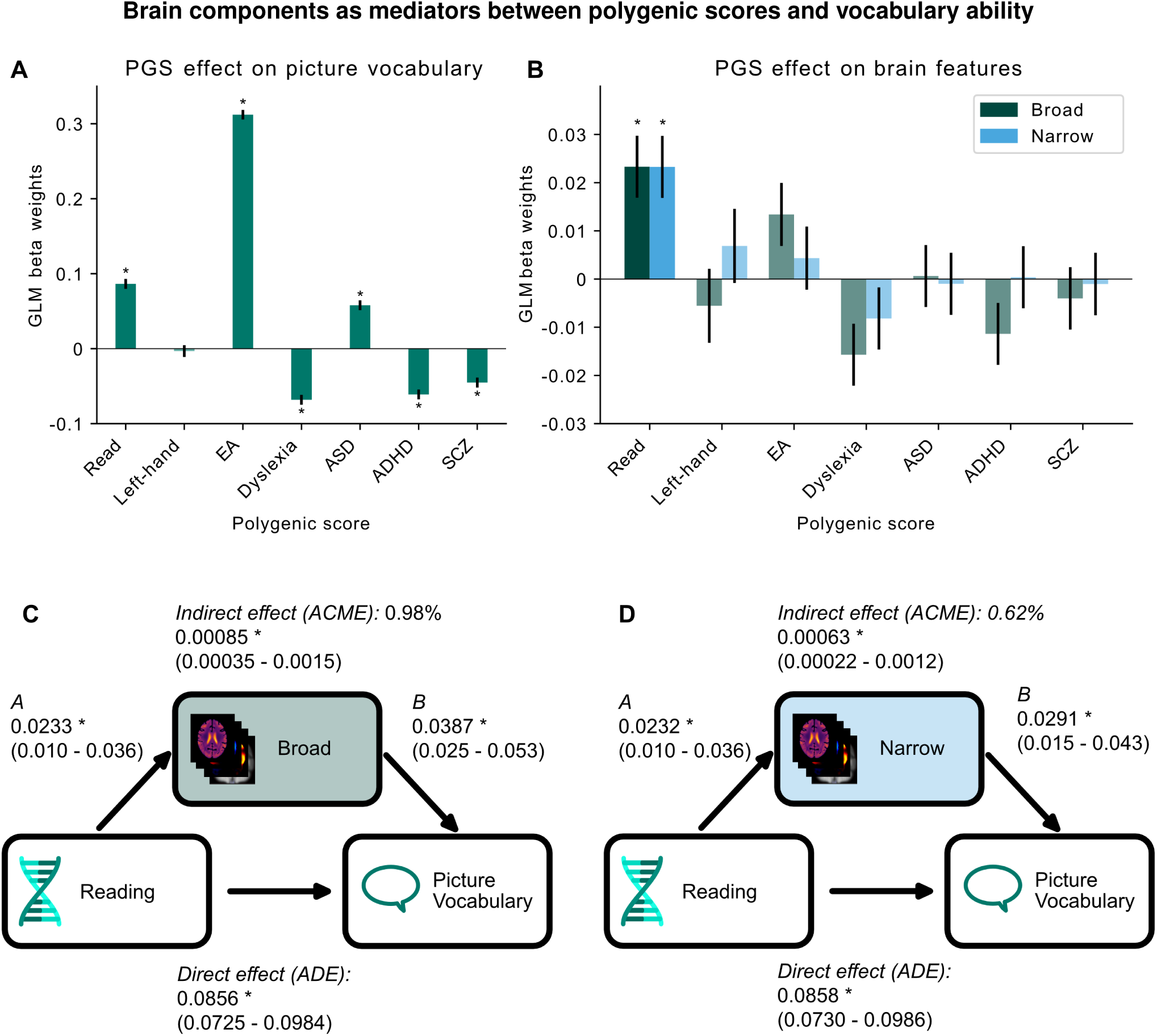
Polygenic score (PGS) and mediation analysis in 21,195 UK Biobank participants. **A.** Effects of PGS for language-related traits (reading ability, left-handedness, educational attainment (EA), dyslexia) and neurodevelopmental conditions (autism spectrum disorder (ASD), attention deficit hyperactivity disorder (ADHD) and schizophrenia (SCZ)) on vocabulary test scores. **B.** Effect of each language-related PGS on brain language components. **C+D.** Mediation analysis of the broad (C) and narrow (D) brain language components, where these components are modeled as mediators of the association between PGS for reading ability and picture vocabulary performance in UK Biobank participants. **A-D.** Asterisks indicate significance after Bonferroni correction for 7 (A), 14 (B), and 10 (C+D) independent tests. **A+B.** Error bars indicate standard error. GLM: generalized linear model **C+D**. A and B indicate beta weights from a generalized linear model with 95% confidence intervals between brackets. ACME: average causal mediation effect. ADE: average direct effect.

### Genomic associations with broad and narrow brain language components identify roles for inhibitory neurons, glia and endothelial cells

For the broad language component we estimated a SNP-based heritability of 29% (GCTA method *h*^2^ = 0.294; LDSC method *h*^2^ = 0.290, see Methods). For the narrow language component we estimated SNP-based heritability at 23-25% (GCTA method *h*^2^ = 0.255, and LDSC method *h*^2^ = 0.232). The two components were substantially, but not completely, genetically correlated (*r_g_* = 0.63, *P* = 4.6 * 10^−66^).

Across both components together we identified 18 genomic loci at genome-wide significance after multiple testing correction for conducting 2 GWAS (Methods). Sixteen loci were associated with the broad structure-function language component (Figure 4, Supplementary Table 4), which mapped to 102 genes (Supplementary Table 5). Eleven loci were associated with the narrow component (Figure 4, Supplementary Table 6), which mapped to 168 genes (Supplementary Table 7). Of the 18 genomic loci, 7 were significantly associated with the broad component alone, 2 significantly with the narrow component alone, and the other 9 significantly with both components. See Supplementary Figure 5 and Supplementary Table 8 for the directions of effect of lead SNPs.

**Figure 4:**
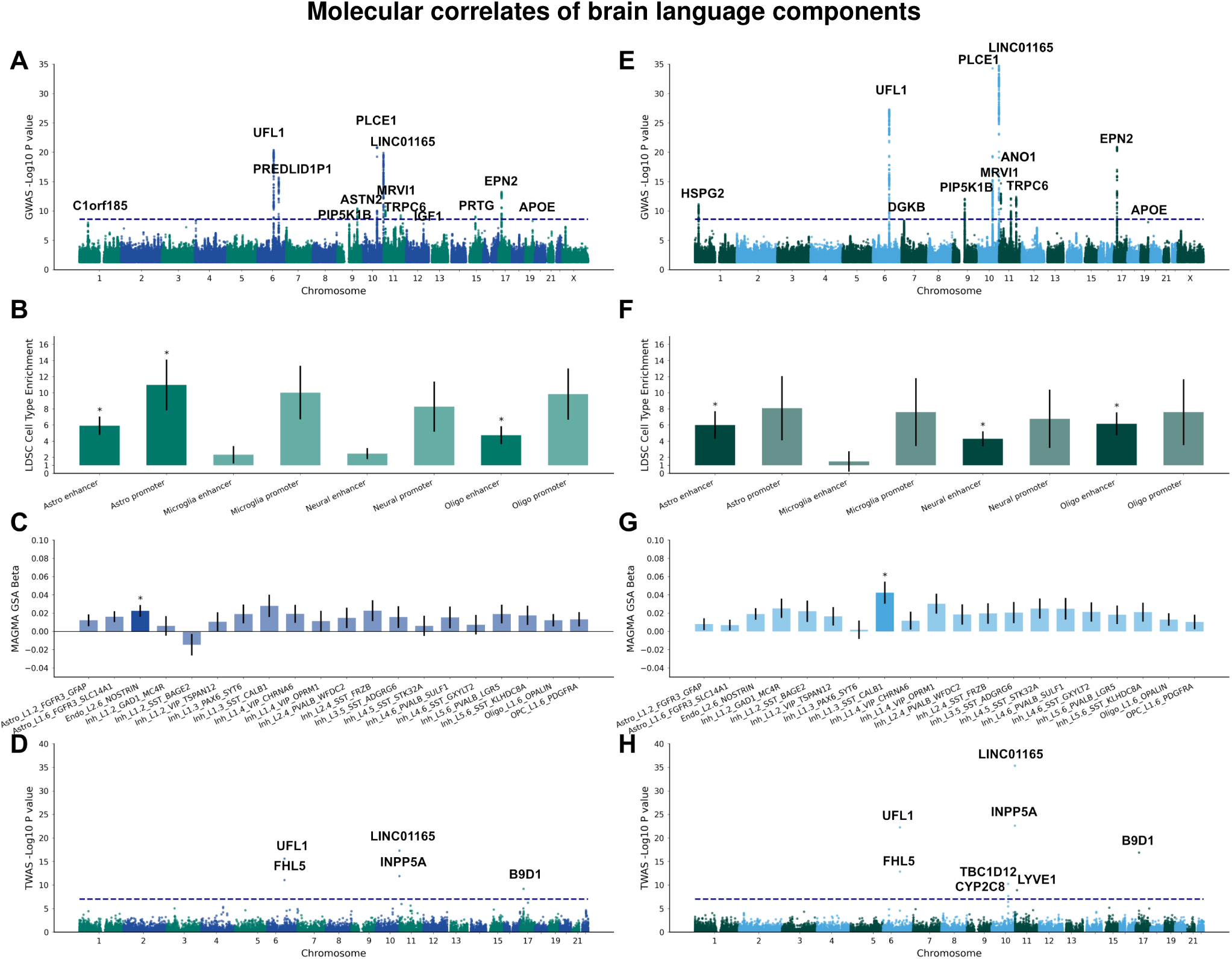
**A +E.** Manhattan plots for GWAS of the broad and narrow structure-function language network components. Dashed lines indicate genome-wide significance (P-value *<* 2.5 * 10^−8^). **B + F.** Cell type enrichment analysis (via LDSC) for GWAS results. **C + G.** Beta weights from gene-set analysis in the superior temporal gyrus in the Allen Human Brain Atlas (via MAGMA). **D + H.** Manhattan plots for TWAS results from S-PrediXcan for inflation-corrected P-values (using Bacon). Only the lowest P-value per gene across tissues is shown. Dashed line indicates Bonferroni correction for number of genes and tissues. **B-F.** Asterisks indicate Bonferroni-corrected significance, black line indicates standard error.

Sixteen of the 18 loci were previously reported in GWAS of other brain image-derived phenotypes in UK Biobank^17,50–53^, with 4 loci previously found in GWAS of language network functional connectivity^23,24^ (see Supplementary Tables 9-10 for a complete list). Two genomic loci were newly identified in the present study to affect human brain variability. The first is located on 11q13.3, with lead SNP rs3929357 (*β_broad_* = 0.044, *P_broad_* = 2.95 * 10^−9^, for T to C, effect allele frequency (EAF): 0.63, Supplementary Figure 6). The nearest gene is *RP11-626H12.1*, which is a long non-coding RNA implicated in blood plasma cancer (multiple myeloma^54^), but not previously in brain phenotypes. The second locus is in 9q21.11 with lead SNP rs2039425 (*β_broad_* = −0.04, *P_broad_* = 1.7 * 10^−8^, for A to G, EAF: 0.65, Supplementary Figure 7), near the gene *PIP5K1B*, encoding a lipid kinase enzyme^55^. The lead SNP is an expression quantitative trait locus of *PIP5K1B* in the cerebellum^56^ (Supplementary Figure 4). Other SNPs near *PIP5K1B* have been associated with general cognitive function^57^ and this gene is also implicated in cerebellar ataxia^58^.

Two of the most significant genomic loci in the present study are also located near genes involved in lipid metabolism and lipid-based signaling: *PLCE1* and *INPP5A*, both located on chromosome 10. The lead SNP near *PLCE1* is rs3891783 (*β_broad_* = 0.069, *P_broad_* = 1.4 * 10^−21^, for C to G, EAF: 0.43, Supplementary Figure 8). For *INPP5A* the lead SNP is rs7084359 (*β_broad_* = 0.067, *P_broad_* = 1.3*^−20^, for A to C, EAF: 0.52, Supplementary Figure 9). *PLCE1* encodes an enzyme that hydrolyzes phospholipids and also acts on the Ras/MAPK-signalling pathway^59^. INPP5A encodes a phosphatase involved in IP3-signalling and is implicated in anterior temporal lobe atrophy^60^.

Partitioned heritability analysis for the broad component showed significant enrichment for astrocyte enhancer and promoter sites, and oligodendrocyte enhancers (Figure 4, Supplementary Table 11). Neuronal cell types and microglia did not show significant enrichment. Cell-type analysis based on gene expression in the superior temporal gyrus from the Allen Human Brain Atlas^61^ showed significant association for endothelial cells (ENDO_L2.6_NOSTRIN, Figure 4, Supplementary Table 12). Gene set analysis using MAGMA showed significant associations of the broad component with supramolecular fibre organization (*P* = 9.5 * 10^−7^) and actin filament organization (*P* = 1.1 * 10^−6^, see Supplementary Table 13). Gene expression across brain development (Brainspan^62^, Supplementary Figure 10) and across tissues (GTEx^56^, Supplementary Figure 11) showed no significant enrichments. Transcriptome-wide association analysis showed the most significant results for *LINC01165* expressed in the cortex, *UFL1* in the caudate, *INPP5A* in the caudate, *FHL5* in the putamen, as well as *B9D1* in the cerebellum (Figure 4, Supplementary Table 14).

Partitioned heritability analysis for the narrow component indicated enrichment for astrocyte, neural and oligodendrocyte enhancer sites (Figure 4, Supplementary Table 15). Cell-type analysis based on gene expression in the MTG from the Allen Human Brain Atlas^61^ showed significant association of inhibitory neurons (Inh_L1.3_SST_CALB1, Figure 4, Supplementary Table 16. Further gene set analysis did not reveal significant associations (Supplementary Table 17). The genetic architecture of the narrow component is significantly associated with nerve tissue in GTEx tissue types^56^ (Supplementary Figure 12), but not significantly enriched during a particular developmental stage (Supplementary Figure 13). Transcriptome-wide associations showed some overlaps with findings for the broad component: LINC01165 expressed in the cortex, *INPP5A* in the caudate, *UFL1* in the amgydala, *B9D1* in the hypothalamus, *FHL5* in the putamen, *TBC1D12* in the spinal cord, and *LYVE1* in the cortex (Figure 4, Supplementary Table 18).

### Brain language components are linked to genomic signatures of evolutionary conservation and selection

Recent advances in comparative genomics and ancient DNA analysis have provided a diverse set of genomic annotations with potential importance for the evolution of distinct human traits, such as language. Here, we queried various such genomic annotations with respect to the genetic architecture of the broad and narrow brain structure-function language components, including ancient selective sweep sites^63^, Neandertal-introgressed genomic fragments^64^, archaic deserts^65^, fetal brain human-gained enhancers (HGEs)^66^, as well as genomic regional conservation or accelerated evolution scores based on comparative genomic analysis across multiple primate species^67^. Ancient selective sweep sites^63^ are parts of the genome that reached fixation in modern humans at an accelerated pace since the split from Neandertals and Denisovans, roughly 650,000 years ago, and before modern humans migrated out of Africa, ~100,000 years ago, and therefore provide possible insights into positive selection for Homo sapiens-specific traits^63^ (Methods). Neandertal-introgressed fragments are regions that were introduced into the Homo sapiens genomic background via archaic admixture events ~50,000 to 60,000 years ago. In contrast, archaic deserts represent genomic regions that are significantly depleted of Neandertal ancestry^65^, which have previously been associated with present-day human variation in inferior frontal cortex structure and/or the uncinate fasciculus fibre tract that links temporal and frontal language regions^68^. Fetal brain HGEs were identified from post-translational histone modification profiles that differ across human, macaque, and mouse^66^ and represent evolutionary changes over the last ~30 million years.

SBayesS analysis (Methods) showed evidence of moderate negative selection for both the broad (*S* = −0.49, *SD*: 0.13), and narrow (*S* = −0.48, *SD*: 0.14) brain language components, which is in line with previous observations for a range of human cognitive traits^67,69^.

Partitioned heritability analysis showed that the SNP-heritability of the broad brain language component is enriched in fetal brain HGEs, and conserved primate phastCons annotations (Figure 5, Supplementary Table 19). MAGMA gene set analysis with key evolutionary gene sets showed no significant associations. (Figure 5, Supplementary Table 20). On the other hand, the narrow language component did not display any significant partitioned heritability enrichment or depletion (Figure 5C, Supplementary Table 21), but MAGMA analysis revealed significant enrichment in ancient selective sweep sites (Figure 5, Supplementary Tables 22-23).

**Figure 5:**
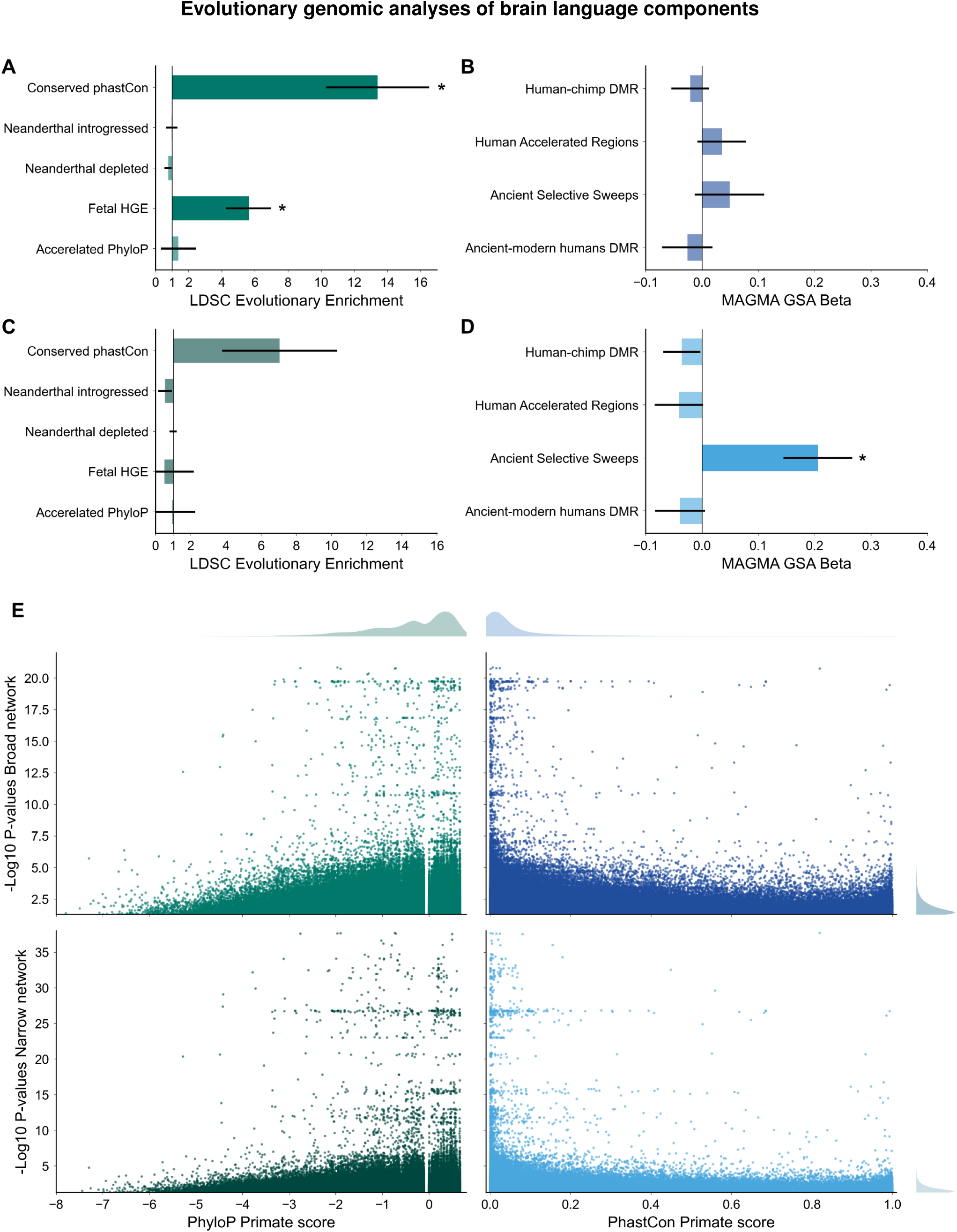
**A+C.** SNP-heritability enrichment/depletion results for evolutionary annotations including accelerated primate phyloP scores, fetal brain human- gained enhancers (HGEs), archaic deserts, Neandertal introgressed fragments, and Conserved primate phastCons scores. **B+D.** Gene set analysis results for differentially methylated regions (DMRs) between anatomically modern and archaic humans (Ancient-modern human DMR), ancient selective sweep sites, Human Accelerated Regions (HARs), and human-chimpanzee differentially methylated regions (Human-chimp DMR). **E.** Scatter plots show the relationship of GWAS −log(*P-values*) of 955,932 nominally significant SNPs associated with either or both of the broad language component (top) and the narrow language component (bottom) with primate clade-derived phyloP scores in the left column, and primate PhastCons scores in the right column. PhyloP scores indicate convergence (negative) and divergence (positive) among primates and PhastCons indicate levels of conservation (with values close to 1 being highly conserved). **A-D.** Asterisks indicate significance after Bonferroni correction for 10 (LDSC) and 8 (MAGMA) independent tests.

The relationship of GWAS P-values to primate clade phyloP and PhastCons scores for the 955,932 nominally trait-associated variants with either or both of the broad and narrow language components highlighted SNPs in both primate-accelerated and conserved loci, with the variants roughly following a Zipfian distribution (Figure 5). However, there was no clear relationship between GWAS significance and the rate of acceleration/conservation among primates.

## Discussion

In this study we identified multimodal brain imaging components that capture functional connectivity of core language regions together with their structural correlates. Through contextualizing the resulting brain maps in terms of cognitive function and neurotransmitter receptor densities, and using individual participant contributions to the structure-function language components as phenotypes for genetic association analyses, our multimodal imaging approach yields various new insights into language biology that have not been apparent from unimodal approaches.

Language areas have been argued to form a distinct module within the brain^3^, although the question remains to what extent can such a module be delineated, and clearly such a module does not exist in isolation from other networks^2,70,71^. In analyses of our broad structure-function language component, we found that increased functional language connectivity was associated with increased motor and cerebellum volume, reduced volume in sensory areas and ventricles, and reduced white matter integrity in the optic radiation. The broad component also showed association of its structural map with activation maps for reward and emotion processing, as well as motor processing. For our narrow structure-funciton language component, in which higher functional connectivity was associated with higher volume in inferior frontal and temporal lobe and decreased volume in the left temporo-parietal junction, there were various cognitive functional annotations beyond language. Overall, these findings suggest co-dependence and integration of language with multiple other functions and networks. The degree of separation of language from other functions in the brain likely reflects the level of granularity with which it is studied.

In terms of molecular distinctiveness, previous post-mortem work identified higher density of serotonergic (5-HT1A), cholinergic (M2) and kainate receptors in cortical language areas compared to primary sensory areas^36,37^. The first of these, 5-HT1A we replicated in correlation analysis with brain-wide PET maps. More globally, we observed positive correlations of these receptor densities with functional connectivity to core language regions, based on brain-wide PET data, similar to other recent work^72^. In contrast, we found our brain volumetric map of the broad component to negatively correlate in particular with serotonergic PET maps. This is explained by the differences between the structural and functional maps, highlighting different areas of the brain, and might indicate a compensatory mechanism related to ageing (whereby declining regional grey matter volume is accompanied by increased serotonergic receptor density).

In terms of individual differences, we found that polygenic disposition for word reading ability was associated with the broad and the narrow structure-function language components. We then found that the broiad and narrow structure-function language components can partly mediate an association between polygenic scores for reading ability and vocabulary levels. Interestingly, the reading polygenic score, based on SNP-wise effect sizes from a GWAS in roughly 27,000 participants^38^, was more strongly associated with our structure-function language components than polygenic scores for dyslexia^42^ or educational attainment^40^, where SNP-wise effect sizes came from GWAS studies with much larger sample sizes (over 700,000 for educational attainment and over one million for dyslexia). Larger GWAS are normally expected to provide less noisy and more predictive polygenic scores. However educational attainment, as measured by the highest education level achieved by participants^40^, and dyslexia as measured by a categorical self-report of having been diagnosed in the GWAS study of Doust et al.^42^, may be less sensitive to individual variation than quantified task performance such as used in the GWAS of reading ability^38^

A potential role of the cerebellum in language is a matter of key interest. Over the past decade evidence in support of this has accumulated based on functional imaging^73^ and lesion studies^74^. Our findings suggest there may be integration between functional language connectivity and structural variation in the cerebellum. However, we also see that this only leads to a modest increase in mediated effects of reading PGS on picture vocabulary, as such effects were similar for the broad component including the cerebellum and the narrow component that did not have a strong cerebellar contribution.

Only 4 of 18 loci identified in the present study overlap with previous unimodal imaging-based GWAS studies of functional connectivity of language networks in the UK Biobank^23,24^, which supports the utility of a multimodal imaging approach as used in the present study. The genetic architecture of the broad structure-function language component provides evidence for roles of astrocytes and oligodendrocytes, as well as endothelium-related gene expression, such that supportive cells in the central nervous system are implicated more strongly than neuronal cells. Given that the broad component has a structural volumetric contribution that was substantial and widespread throughout the brain, tissue growth and maintenance may be especially important, reflected in key contributions from supportive cells. This may also relate to the enrichment for primate conserved genetic elements and fetal human-gained enhancers that we observed. Somewhat differently, for the narrow component, the genetic architecture provides evidence for the role of neurons, especially cortical inhibitory neurons, as well as astrocytes and oligodendrocytes, and genes near ancient selective sweep sites. The latter findings would suggest that a degree of tuning of inhibitory neurophysiology and supportive cells to support language may have occurred relatively recently in human evolution, since our split from Neandertals^75,76^.

The multimodal imaging approach of this study helped us identify structural variation outside of canonical language areas that is nonetheless linked to core language region connectivity. In fact, tensor-based morphometry captures larger overall variance in the brain than white matter integrity (apparent fibre density) and ICA resting state maps. Furthermore, by focusing only on the resting functional connectivity of inferior frontal and superior temporal language-related regions, we only enter a small fraction of the brain’s overall resting state signal into the linked-ICA decomposition, but all of the variance in brain structure. These factors likely contributed to the general dominance of TBM for most of the multimodal components derived from linked-ICA. In addition, the variance within modalities is larger than that shared across modalities, such that there were few multimodal components identified by the linked-ICA even when deriving relatively small numbers of components overall. Nonetheless, the 5c and 10c decompositions produced one component each that was sufficiently multimodal to enrich biological discovery in relation to language organization in the brain.

In this study we integrated brain structural and functional metrics to derive multimodal components that capture language network connectivity and its structural correlates in a large adult dataset. This approach sheds light on brain structure-function relations and provides novel phenotypes for genetic discovery. We see that brain language connectivity is related to structural variation beyond core language areas, including notably the cerebellum, and that 23-30% of the variance in the structure-funciton language components is explained by common genetic variation. We identify 2 genomic loci newly associated with human brain traits, 14 new genomic loci for brain language organisation, and confirm the importance of 4 other loci previously associated with brain language organisation. Both neuronal and non-neuronal cell types are implicated, as well as evolutionarily conserved and divergent genetic loci, which suggests complex interplays rather than simple scenarios of language emergence or development. This work provides a basis for future mechanistic insights into how genetic factors influence both brain structure and function to shape our language abilities. Future work could include additional imaging modalities in the linked ICA framework, as well as testing the resulting brain components in relation to rare genetic variants with possibly larger effects in their carriers.

## Methods

### Participants

As described previously^23^, imaging and genomic data were obtained from the UK Biobank^77^ as part of research application 16066 from primary applicant Clyde Francks. The UK Biobank received ethical approval from the National Research Ethics Service Committee North West-Haydock (reference 11/NW/0382), and all of their procedures were performed in accordance with the World Medical Association guidelines. Informed consent was obtained for all participants^78^. Analyses were conducted on 32,677 participants that remained after quality control of genotype and imaging data (see below).

### Imaging data

Brain imaging data were collected for UK biobank as described previously^23,33,79^. Identical scanners and software platforms were used for data collection at different sites (Siemens 3T Skyra; software platform VD13). For collection of rs-fMRI data, participants were instructed to lie still and relaxed with their eyes fixed on a crosshair for a duration of 6 minutes. In that timeframe 490 datapoints were collected using a multiband 8 gradient echo EPI sequence with a flip angle of 52 degrees, resulting in a TR of 0.735 s with a resolution of 2.4×2.4×2.4mm^3^ and field-of-view of 88×88×64 voxels. Our study made use of pre-processed image data generated by an image-processing pipeline developed and run on behalf of UK Biobank.

### Genetic data

As described previously^23^, genome-wide genotype data (UK Biobank data category 263) were obtained by the UK Biobank using two different genotyping arrays (for full details see^77^). Imputed array-based genotype data contained over 90 million SNPs and short insertion-deletions with their coordinates reported in human reference genome assembly GRCh37 (hg19). In downstream analyses we used both the non-imputed and imputed array-based genotype data in different steps (below).

### Sample-level quality control

Similar to previous work^23^, sample-level quality control at the phenotypic and genetic level was conducted on 49,179 participants who had imaging and genotype data available. In phenotype sample-level quality control, participants were first excluded with imaging data labelled as unusable by UK Biobank quality control. Second, participants were removed based on outliers (here defined as 3 standard deviations from the mean) in at least one of the following metrics: discrepancy between rs-fMRI brain image and T1 structural brain image (UK Biobank field 25739), inverted temporal signal-to-noise ratio in preprocessed and artefact-cleaned preprocessed rs-fMRI (data fields 25743 and 25744), scanner X, Y and Z brain position (fields 25756, 25757 and 25758) or in deviation from template (see section Imaging data preprocessing and phenotype derivation). In total 8,282 participants were excluded in the phenotype QC.

Subsequently, in genetic sample-level quality control, only participants in the pre-defined European ancestry cluster were included (data-field 22006)^77^, as this was the largest single cluster in terms of ancestral homogeneity – an important consideration for some of the genetic analyses that we carried out (below). Furthermore, participants were excluded when self-reported sex (data-field 31) did not match genetically inferred sex based on genotype data (data-field 22001), or when exclusion thresholds were exceeded in heterozygosity (≥ 0.1903) and/or genotype missingness rate (≥ 0.05) (data-field 22027). Finally, one random member of each pair of related participants (up to third degree, kinship coefficient ≥ 0.0442, pre-calculated by UK Biobank) was removed from the analysis. This led to the further exclusion of 8,220 participants. In total 32,677 participants were included in all further analyses.

### Imaging data pre-processing and phenotype derivation

Imaging phenotype derivation consisted of two main steps. First, we derived unimodal brain maps from different MRI imaging modalities, which were then entered together in a dimensionality reduction technique (linked-ICA) to derive data-driven independent components that can share variation across imaging modalities.

#### Structural MRI: tensor-based morphometry

Using the minimally processed T1-weighted MRI scans from UK Biobank^33,79^, tensor-based morphometry (TBM) maps were derived as described previously^31^. In brief, a study-specific template was generated using from a subset of 1000 randomly chosen individuals, using Advanced Normalization Tools (ANTs v. 2.3.5)^80^. Symmetric image normalization (SyN) was used to register each subject’s T1-weighted image non-linearly to this template, after being histogram-matched and winsored at 1-99 percentiles. Total field variance was set to 3, and update variance was set to 0, with a downsampling scheme of 6-4-2-1 and smoothing sigmas of 4,2,1 and 0 voxels. The affine registration matrix and deformation fields were converted to the Jacobian determinant maps, a measurement of local tissue deformation compared to the study template. 2,098 individuals that failed ANTs affine registration were instead registered using a similar approach, FSL Flirt^81^. This processing difference was included as a binary covariate for further analyses. Jacobian determinant deformation maps were used as IDP maps into multimodal decomposition.

#### Diffusion MRI: fixel-based analysis

Fixel-based maps were derived as described previously^31^. In brief, for our analysis of white matter integrity we used minimally-preprocessed dMRI scans from UK Biobank^33,79^, with a 2mm^3^ spatial resolution. Diffusion data were acquired with b-values of 1,000 and 2000 s/mm^2^ using a multiband acceleration factor of 3. For both b-values, the diffusion-weighted shell was covered with 50 distinct diffusion-encoding directions. Minimal preprocessing by the UK Biobank team consisted of correction for off-resonance warps, eddy currents, head-motion and gradient nonlinearity. These corrections were rerun for 8,245 individuals^31^, controlling for this using a binary batch covariate. The main diffusion processing pipeline was run using MRTrix3 v3.0.3^82^, first by constructing a study-specific fiber orientation density (FOD) template based on 890 subjects with good quality data after visual inspection of 1,000 subjects. The template was constructed using in order: on the diffusion data of each subject a N4 bias field correction and intensity normalization, and estimation of the average tract response function^32^. These estimated response functions were used in spherical deconvolution to generate individual whole-brain FOD volumes, which were then registered in a non-linear fashion to a common template and iteratively yielded an average FOD template. To identify the main directions of white-matter tracts in each voxel, fixels were estimated on the FOD template. These processing steps were applied to all subjects afterwards, which were registered to the study-specific template. Each individual FOD volume was segmented, resulting in fixel-based readouts that were transformed to template space^82^. Quality control consisted of visual inspection of all template-transformed zeroth-order harmonic maps, which represents average diffusion in each voxel. Apparent fiber density can be derived from the FOD, which is suggested to be approximately proportional to axon fibre bundle density in the primary direction in that fixel^32,83^. AFD maps were used as IDP maps for multimodal decomposition.

#### Resting state MRI: language-related connectivity maps

Preprocessing was conducted by the UK Biobank and consisted of motion correction using MCFlirt^84^, intensity normalization, high-pass filtering to remove temporal drift (sigma=50.0s), unwarping using fieldmaps and gradient distortion correction. Structured scanner and movement artefacts were removed using ICA-FIX^85–87^. Preprocessed data were registered to a common reference template in order to make analyses comparable (the 6th generation nonlinear MNI152 space, http://www.bic.mni.mcgill.ca/ServicesAtlases/ICBM152NLin6) and smoothed with a Gaussian filter (*σ* = 2.5) using FSL^81^. To obtain resting state language network maps, we regressed each subject’s time courses from the previously derived group-ICA components (100 component decomposition)^79^ on each subject’s resting state fMRI data, which is equivalent to the second step of dual regression^81^. The generated connectivity maps for component 28 (left inferior frontal gyrus) and 9 (left temporal lobe), roughly overlapping with core nodes of the language network, were used as IDP maps for multimodal decomposition.

#### Dimensionality reduction with linked independent component analysis

After genetic and imaging QC and preprocessing, 32,677 participants had all 4 unimodal IDP maps available. These were entered into a linked-ICA decomposition with 5 or 10 components. Linked-ICA is a multimodal extension of ICA, using a shared mixing matrix^10,27^. This allows for the derivation of shared patterns across multiple imaging modalities. Linked-ICA provides, for each component, spatial maps for each contributing imaging modality, as well as participant courses that indicate the contribution of each participant to the component, and modality loadings, indicating the relative contribution of each modality to that component. For each component we calculated a multimodal index (MMI), as previously described^88^. For the purposes of this study we sought multimodal components with > 0.4 on MMI, a minimum resting state contribution > 0.2 on unimodal resting state maps, and low individual participant dominance (< 0.01).

We corroborated model order selection by looking at the knee of the scree plots of unimodal principal component analysis plots (Supplementary Figure 3), which suggested between 5 and 10 as an optimal range for the number components for a multimodal ICA decomposition. To check that these relatively low numbers of components did not unduly affect the outcome, we also ran a 50-component decomposition (Supplementary Figure 14), which showed very similar phenotypic and genetic profiles for the resting state dominant components (Supplementary Figures 15-16).

### Post-image analysis

To understand the cognitive and neurotransmitter architectures of our multimodal brain language components, we decoded them using NiMARE^28^ CorrelationDecoder in the Neurosynth LDA-50 (version 7)^29^ and the PET neurotransmitter maps from the neuromaps databases^30^. Note that due to the fixel-based data structure the AFD maps could not be decoded using NiMARE.

### Genetic variant-level QC

Similar to previous work^23^, three different genetic datasets were prepared, as needed for four different analysis processes:

1. Array-based genotype data were filtered, maintaining variants with linkage disequilibrium (LD) ≤ 0.9, minor allele frequency (MAF) ≥ 0.01, Hardy-Weinberg Equilibrium test *P*-value ≥ 1 × 10^−15^ (see^89^), and genotype missingness ≤ 0.01 for REGENIE step 1 (below).
2. Imputed genotype data were filtered, maintaining bi-allelic variants with an imputation quality ≥ 0.7, Hardy-Weinberg Equilibrium test *P*-value ≥ 1 × 10^−7^ and genotype missingness ≥ 0.05 for association testing in REGENIE step 2.
3. For genetic relationship matrices, SNPs were only used if they were bi-allelic, had a genotype missingness rate ≤ 0.02, a Hardy Weinberg Equilibrium *P*-value ≥ 1 × 10^−6^, an imputation INFO score ≥ 0.9, a MAF ≥ 0.01, and a MAF difference ≤ 0.2 between the imaging subset and the whole UK Biobank.

### Heritability and genetic correlation analysis

As described previously^23^, genetic relationship matrices (GRMs) were computed for the study sample using GCTA v. 1.93.0beta^90^. In addition to the previous sample-level quality control, individuals with a genotyping rate ≤ 0.98 and one random individual per pair with a kinship coefficient ≥ 0.025 derived from the GRM were excluded from heritability analysis. The SNP-based heritability of both components was estimated using genome-based restricted maximum likelihood (GREML) in GCTA v. 1.93.0beta^90^. Heritability was also quantified using linkage disequilibrium score regression (LDSC v1.0.1) (based on the European subset of the 1000 Genomes Phase 3 reference panel) as a sensitivity analysis and also because this pipeline supported downstream analysis using partitioned heritability. To investigate the genetic similarity between the broad and narrow language components, we also used LD score regression v.1.0.1^91^ to compute the genetic correlation.

### Mediation analysis

We tested (1) the association between polygenic scores for reading ability and picture vocabulary scores (UK Biobank field 26302-2-0), (2) the association between polygenic scores for reading ability and brain language components and (3) to what extent the brain language components mediate the association of polygenic scores for reading ability with picture vocabulary scores. The picture vocabulary test in UK Biobank is a computer adaptive test to measure a participant’s vocabulary level, adapted from the NIH Toolbox^92^ and recalibrated for use in a UK cohort^93^. All of the words within the test are rated with a difficulty score from −11 to 11. In each round, the participant is shown a word on a screen and has to choose the correct picture for that word. If the picture is chosen correctly, the participant will get a more difficult word next round, if not, the next word will be easier. Picture vocabulary is a reliable indicator of linguistic knowledge^94^. The picture vocabulary test is the only test of a linguistic cognitive function available in UK Biobank for more than 5000 participants.

Polygenic scores in UK Biobank individuals were calculated based on SNP-wise effect sizes from previous GWAS studies: for reading abilities (quantitatively assessed in up to 33,959 participants from the GenLang consortium)^38^, dyslexia (51,800 cases and 1,087,070 controls) from 23andMe Research Institute^42^ and left-handedness (UK Biobank participants without imaging data: 33,704 cases and 272,673 controls)^39^, educational attainment (765,283 individuals), autism spectrum disorder (18,381 cases, 27,969 controls), attention deficit hyperactivity disorder (38,691 cases, 186,843 controls) and schizophrenia (76,755 cases, 243,649 controls). We derived polygenic scores with PRS-CS 51 and used a generalized linear model and mediation analysis (using the package mediation^95,96^) in R (v4.40) to test these questions. PRS-CS^97^, a Bayesian regression framework to infer posterior effect sizes of autosomal SNPs based on genome-wide association summary statistics, was applied using default parameters and a recommended global shrinkage parameter phi = 0.01, combined with LD information from the 1000 Genomes Project phase 3 European-descent reference panel. PRS-CS performed in a similar way to other polygenic scoring methods, with noticeably better out-of-sample prediction than a clumping and thresholding approach^98,99^. Covariates included in the model were genetic sex, age, age^2^, age × sex, the first 10 genetic principal components that capture genome-wide ancestral diversity, genotype array (binary variable) and various scanner-related quality measures (Supplementary Table 24). 21,195 participants with both brain component scores and picture vocabulary scores available were included in this analysis.

### Genome-wide association testing

Similar to previous work^23^, REGENIE v.3.6.0 was used^89^. In brief, REGENIE is a two-step machine learning method that fits a whole genome regression model and uses a block-based approach for computational efficiency. In REGENIE step 1, array-based genotype data were used to estimate the polygenic signal in blocks across the genome with a two-level ridge regression cross-validation approach. The estimated predictors were combined into a single predictor, which was then decomposed into 23 per-chromosome predictors using a leave one chromosome out (LOCO) approach, using a block size of 1000, 4 threads and low-memory flag. These LOCO polygenic predictors were used as covariates in both common and rare variant association testing in REGENIE step 2. Phenotypes were transformed to a normal distribution in both REGENIE step 1 and 2. Covariates for both steps included genetic sex, age, age2̂, agesex, the first 10 genetic principal components that capture genome-wide ancestral diversity^77^, genotype array (binary variable) and various scanner-related quality measures (scanner X, Y and Z-position, inverted temporal signal to noise ratio and mean displacement as an indication of head motion, amount of T1 warping to standard space, discrepancy between rfMRI brain image and T1, discrepancy between dMRI and T1, number of dMRI outlier slices, and eddy standard deviations).

In REGENIE step 2, we performed separate single-variant association analyses with common (imputed) variants separately for the broad and narrow brain language components using a Firth regression framework. GWAS were run on our local compute cluster using a block size of 500, 4 threads and a low-memory flag.

Genome-wide significant variants, Bonferroni-adjusted for two GWASs (i.e. threshold *P* = 2.5 * 10^−8^), were annotated using the online FUMA platform (version 1.5.2)^100^. MAGMA (version 1.08)^101^ was used to calculate gene-based *p*-values and to investigate potential gene sets of interest^102,103^ and to map the expression of associated genes in a tissue-specific^56^ and time-specific^62^ fashion. Gene sets smaller than 10 were excluded from the analysis. We also conducted the first and second step of the cell-type analysis implemented in FUMA^104^ on the Allen Brain Atlas human superior temporal gyrus dataset^61^. The conditional results (second step) were reported.

### Cell-type analysis using partitioned heritability

To investigate the role of different cell-types in relation to the brain structure-function language components, we applied eight human genome annotations developed by Nott et al.^67,105^ tagging promoter and enhancer regions of neurons, oligodendrocytes, microglia, and astrocytes within LDSC partitioned heritability analysis^91,106^. All enrichment analyses were controlled for the baselineLD model v2.2. Enrichment P-value results were Bonferroni-corrected for 16 tests.

### Transcriptome-wide association study

We ran a TWAS using S-PrediXcan framework^107^ and the joint-tissue imputation (JTI) TWAS derived models from GTEx v8 tissues^108^. S-PrediXcan is an extension to GWAS summary statistics^107^ of the PrediXcan methodology which estimates gene expression from the genotype profile of each individual by using the JTI model weights, which were trained on GTEx^109^, and validated on PsychEncode^110^ and GEUVADIS^111^. S-PrediXcan and JTI weights account for LD and collinearity problems owing to high expression correlation across tissues^107^. We further corrected for genomic inflation using the Bacon package^112^ and corrected with a Bonferroni correction for 531388 tests, accounting for number of tissues, genes and phenotypes (with an associated P-value of 9.4 * 10^−8^).

### Genome-wide negative selection estimation

Similar to previous work^67^, we performed SBayesS analysis on both sets of GWAS summary statistics using the GCTB software (version 2.02)^69^ to quantify the level of genome-wide negative selection. SBayesS estimates total SNP-heritability, polygenicity, and the relationship between variant minor allele frequencies and effect sizes, and generates a genome-wide negative selection metric (*S*), which ranges from 0 to −1. *S* estimates that are closer to −1 are interpreted as a sign of strong negative selection, whereas estimates closer to 0 can suggest positive selection.

### Evolutionary genomic analyses

We employed LD Score regression (LDSC v1.0.1)^91^ to quantify partitioned SNP heritability enrichments or depletions across multiple evolutionary annotations, with particular emphasis on signatures of human-specific evolution and archaic hominin introgression. All analyses were performed against the baselineLD model (v2.2) to account for underlying genomic architecture, provided by the Alkes Price group^106^. Fetal brain human-gained enhancers were also controlled for fetal brain active regulatory elements from the Roadmap Epigenomics Consortium database^113^.

Our evolutionary annotations encompassed:

1. Fetal brain human-gained enhancers (HGEs)^66^. HGEs were previously defined across three consecutive developmental periods in human fetal cortical tissue^66,68^ compared to mice and marmosets.
2. Neandertal introgressed alleles^64^. We used a previously derived annotation that includes both Neandertal introgressed SNPs, and those in perfect linkage disequilibrium (r2̂=̂ 1), reflecting a relatively recent (3̃5,000-50,000 years ago) evolutionary event (i.e., Neandertal-Homo sapiens admixture).
3. Archaic deserts^65^. We used previously defined archaic deserts that represent genomic regions with significant depletion of Neandertal sequence (average introgression percent < 10^−3.5^) across Europeans, East Asians, South Asians, and Melanesians. From the six identified desert regions spanning 85.3 Mb, we eliminated 128 Neandertal introgressed SNPs (constituting approximately 10^−4^ of the total desert length) along with haplotypes in substantial LD (*r*^2^ *>* 0.6).
4. Primate clade acceleration/conservation signatures. We examined loci conserved throughout primate evolution (Conserved_Primate_phastCons46way annotation from baselineLD), and genomic regions with primate phyloP scores^114^ below −2, potentially indicating accelerated evolution.

For complementary gene-based analyses, we used four additional evolutionary genomic annotations covering evolutionary processes on lineages leading to modern humans from 8̃ million years ago to 3̃5,000 years ago, which were not suitable for LDSC partitioned heritability analysis due to limited SNP-coverage^68^:

1. Ancient selective sweep^63^ sites are parts of the genome where Neandertals and Denisovans are not within the range of human variation, reached fixation at a faster rate than expected under standard evolutionary assumptions, and can therefore provide insight into uniquely *Homo sapiens* traits.
2. Human accelerated regions^115–118^ are regions in the genome that are conserved among vertebrates, but show accelerated evolution on the human lineage.
3. Differentially methylated regions between anatomically modern humans and archaic hominins (AMH-derived DMRs)^119^
4. Differentially methylated regions between humans and chimpanzees (Human-chimp DMRs)^119^

Since these four annotations tag regulatory or selective sweep sites, and individually cover less than 1% of well-imputed SNPs in the 1000 Genomes Phase 3 reference panel, we adapted our methodological approach for their analysis. We constructed gene sets by identifying protein-coding genes (based on NCBI hg19 genome annotation^120^) located within ±1 kilobase of each annotated locus. These gene sets were subsequently analyzed using MAGMA^101^ gene set analysis.

## Data Availability

The primary data used in this study are from the UK Biobank. These data can be provided by UK Biobank pending scientific review and a completed material transfer agreement. Requests for the data should be submitted to the UK Biobank: https://www.ukbiobank.ac.uk. Specific UK Biobank data field codes are given in the Methods section. Dyslexia GWAS summary statistics must be obtained by request to 23andMe Research Institute, via https://research.23andme.com/dataset-access/. Other publicly available data sources and applications are cited in the Methods section. GWAS summary statistics from the present study are made available online within the GWAS catalog: https://ebi.ac.uk/gwas/.

## Code Availability

This study used openly available software and codes, specifically: REGENIE (https://rgcgithub.github.io/regenie/), GCTA (https://cnsgenomics.com/software/gcta/#GREML^90^), FUMA (https://fuma.ctglab.nl/), MAGMA (https://ctg.cncr.nl/software/magma), PRS-CS (https://github.com/getian107/PRScs), LDSC (https://github.com/bulik/ldsc), FLICA (https://fsl.fmrib.ox.ac.uk/fsl/docs/utilities/flica.html), ANTs(https://github.com/ANTsX/ANTs), mrtrix (https://mrtrix.readthedocs.io/en/latest/index.html), FSL (https://fsl.fmrib.ox.ac.uk/fsl/docs/index.html, S-PrediXcan (https://github.com/hakyimlab/MetaXcan), NiMARE (https://nimare.readthedocs.io/en/stable/), neuromaps (https://neuromaps-main.readthedocs.io/en/latest/), PLINK (https://www.cog-genomics.org/plink2/), and GCTB (http://www.cnsgenomics.com/software/gctb/). Custom code is available on Github (https://github.com/jamelink/structure_function_language).

## Acknowledgements

This research was funded by the Max Planck Society (Germany). Brain imaging, genomic and cognitive data were obtained from the UK Biobank as part of research application 16066 from primary applicant Clyde Francks. Our study partly made use of pre-processed image data generated by an image-processing pipeline developed and run on behalf of UK Biobank. The funders had no role in study design, data collection and analysis, and the decision to publish or preparation of the manuscript. The authors would like to thank the participants of UK Biobank. The authors would like to thank the research participants and employees of 23andMe Research Institute for making this work possible. The authors would like to thank Simon de Vries for help with figure design choices.

## Author Contributions

Conceptualization - J.S.A., C.F.B., K.V.H., S.E.F., C.F.; Methodology - J.S.A., S.S-N., G.A., M-Y. W., A.L., C.F.B., K.V.H., D.S.; Software - J.S.A., S.S-N., G.A., M-Y. W., A.L., D.S.; Formal analysis - J.S.A., S.S-N.; Data curation - J.S.A.; Writing - original draft - J.S.A.; Writing - review & editing - S.S-N., G.A, D.S., M-Y W., K.V.H., S.E.F, C.F.B., C.F; Visualization - J.S.A.; Project administration - C.F.; Resources - S.E.F, C.F.; Funding acquisition - C.F., S.E.F.; Supervision - S.E.F., C.F.B., C.F.,

## Supporting Information Appendix (SI)

All supplementary figures and tables can be found in accompanying PDF (figures) and Excel (tables/supplementary data) files. Genome-wide multivariate summary statistics from our GWAS based on common genetic variants are available online within the GWAS catalog (https://ebi.ac.uk/gwas/).

