## Supplementary Figures for "Integrating brain structure and function for the neurobiology and genetics of language"

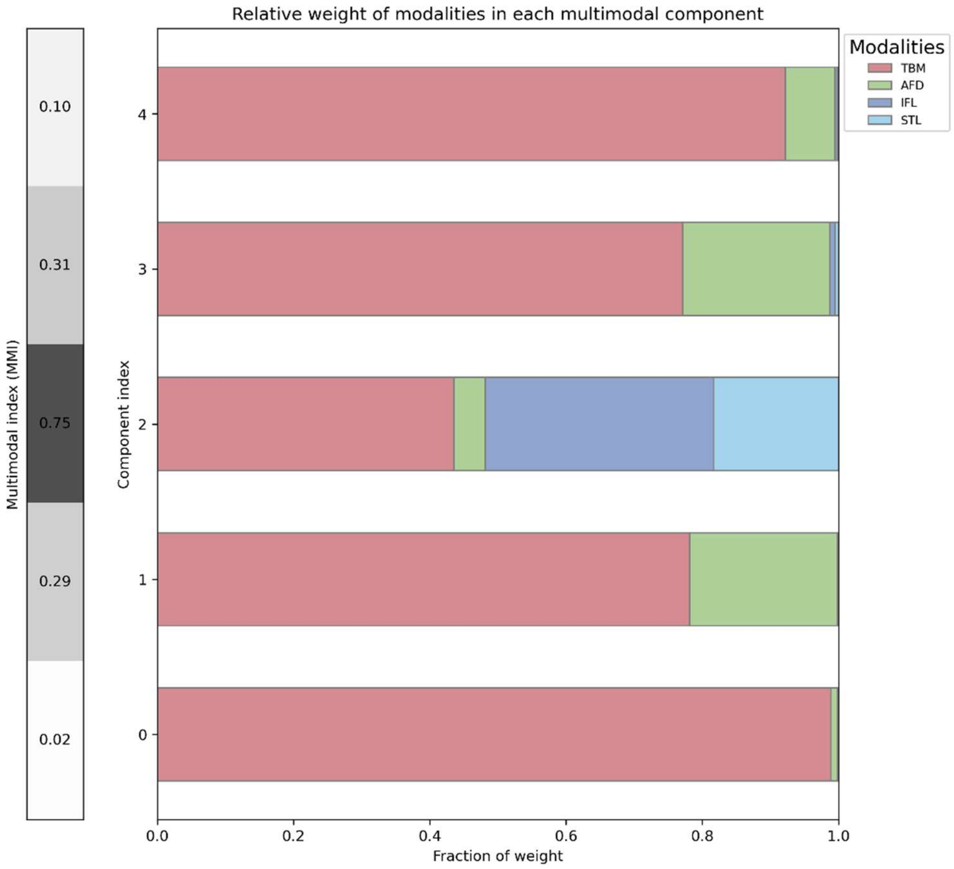


**Supplementary Figure 1.** Fraction weights per modality for all components in the 5 component decomposition. On the left the multimodal index (MMI) is shown, indicative of multimodality. Component 2 was used in the main analysis as the broad brain language network component.

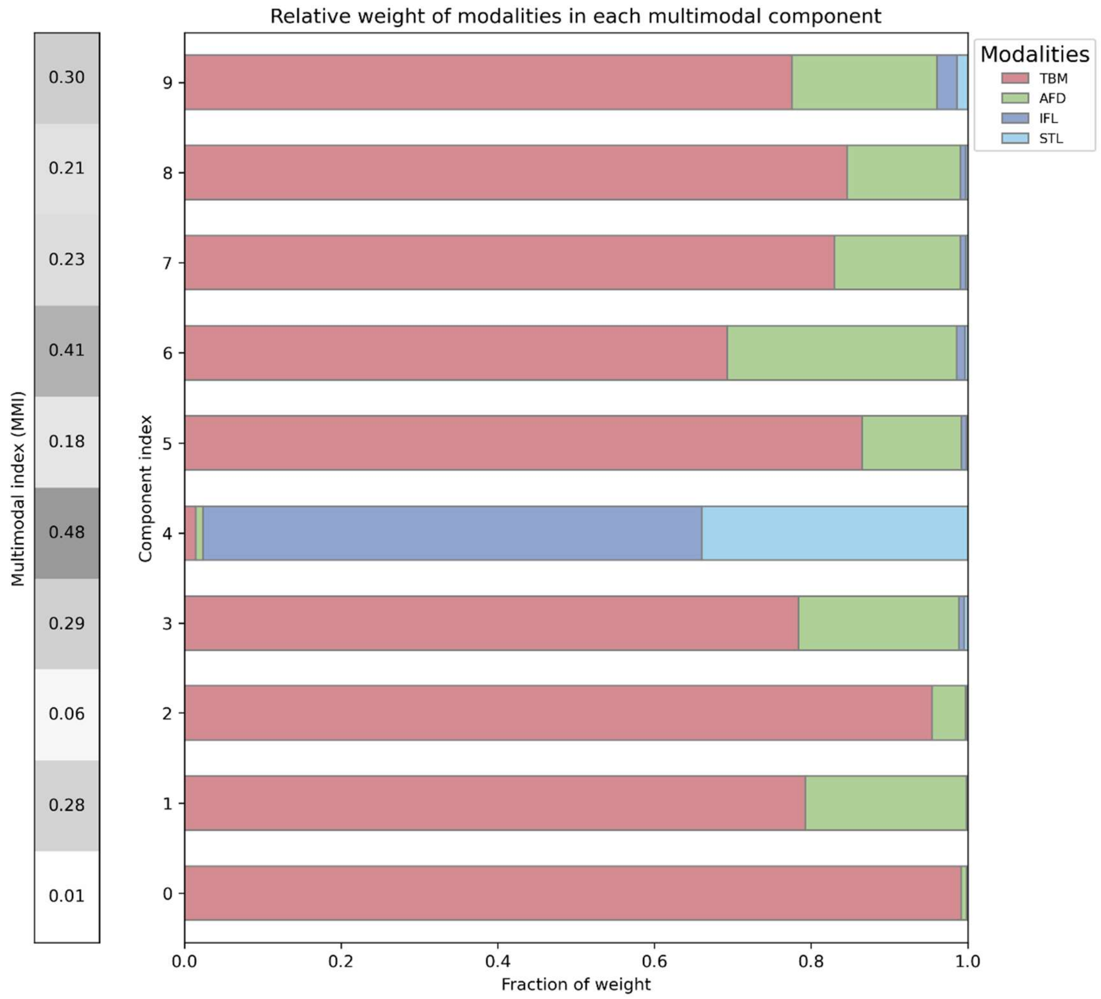


**Supplementary Figure 2.** Fraction weights per modality for all components in the 5 component decomposition. On the left the multimodal index (MMI) is shown, indicative of multimodality. Component 4 was used in the main analysis as the narrow brain language network component.

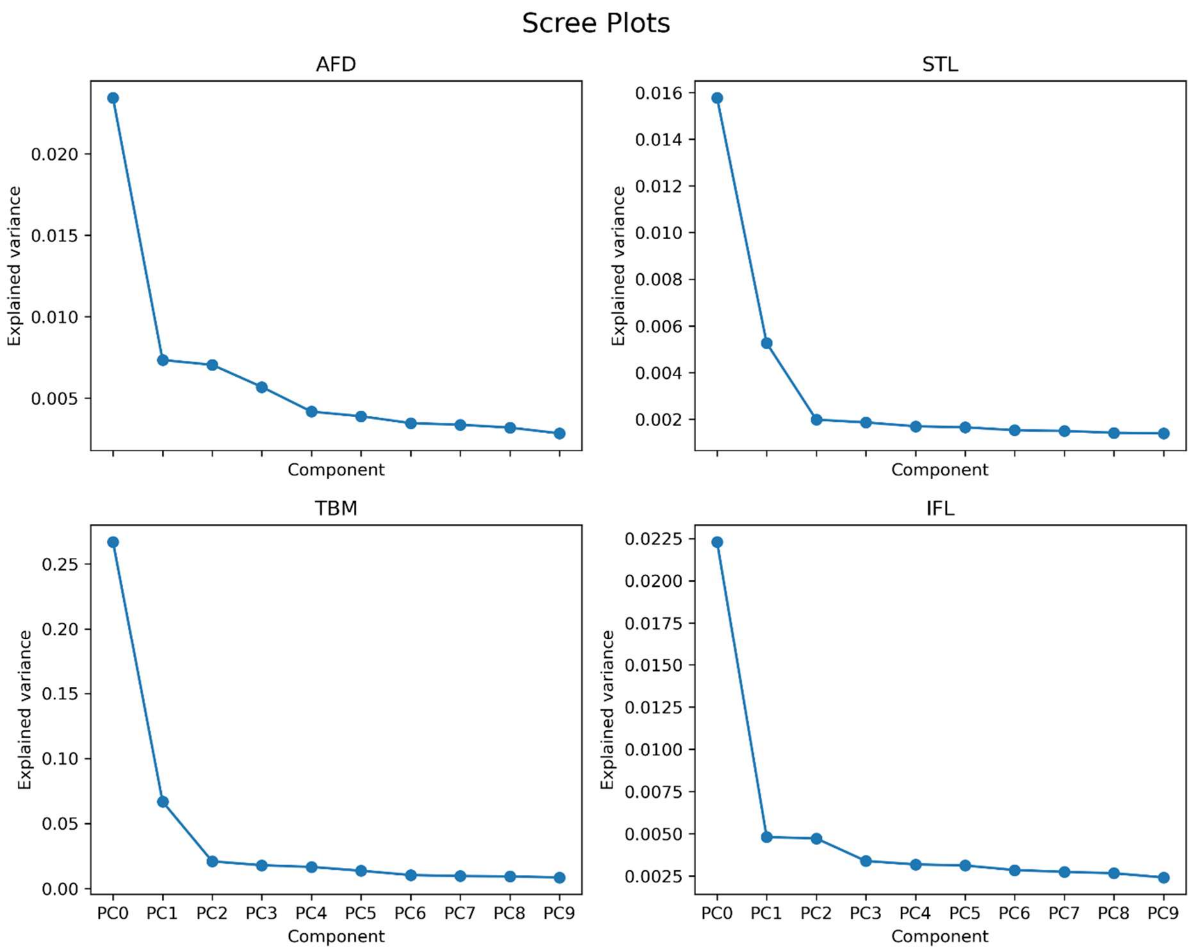


**Supplementary Figure 3.** Explained variance of unimodal PCA decompositions. Elbow points are observed at the first component for AFD and IFL and at the second component for STL and TBM.

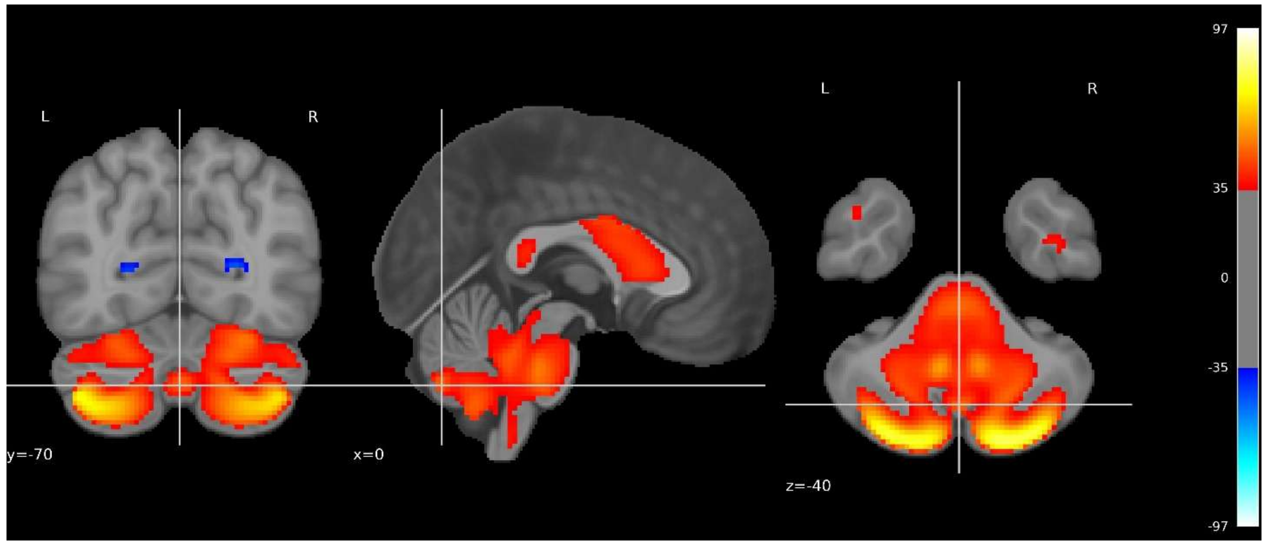


**Supplementary Figure 4 -** Close up figure of the cerebellum in the brain volumetric map of the broad language component.

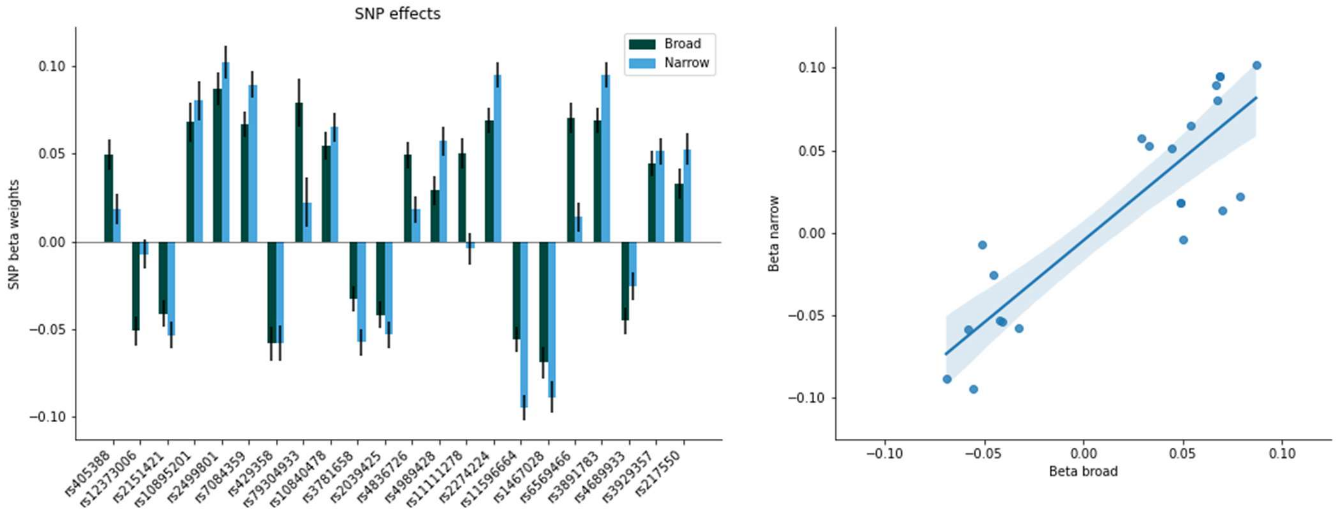


**Supplementary Figure 5.** Directionality of associations of lead SNPs in either GWAS (22 in total). On the left beta weights in a bar plot, on the right the betas plotted against each other for the broad and narrow components.

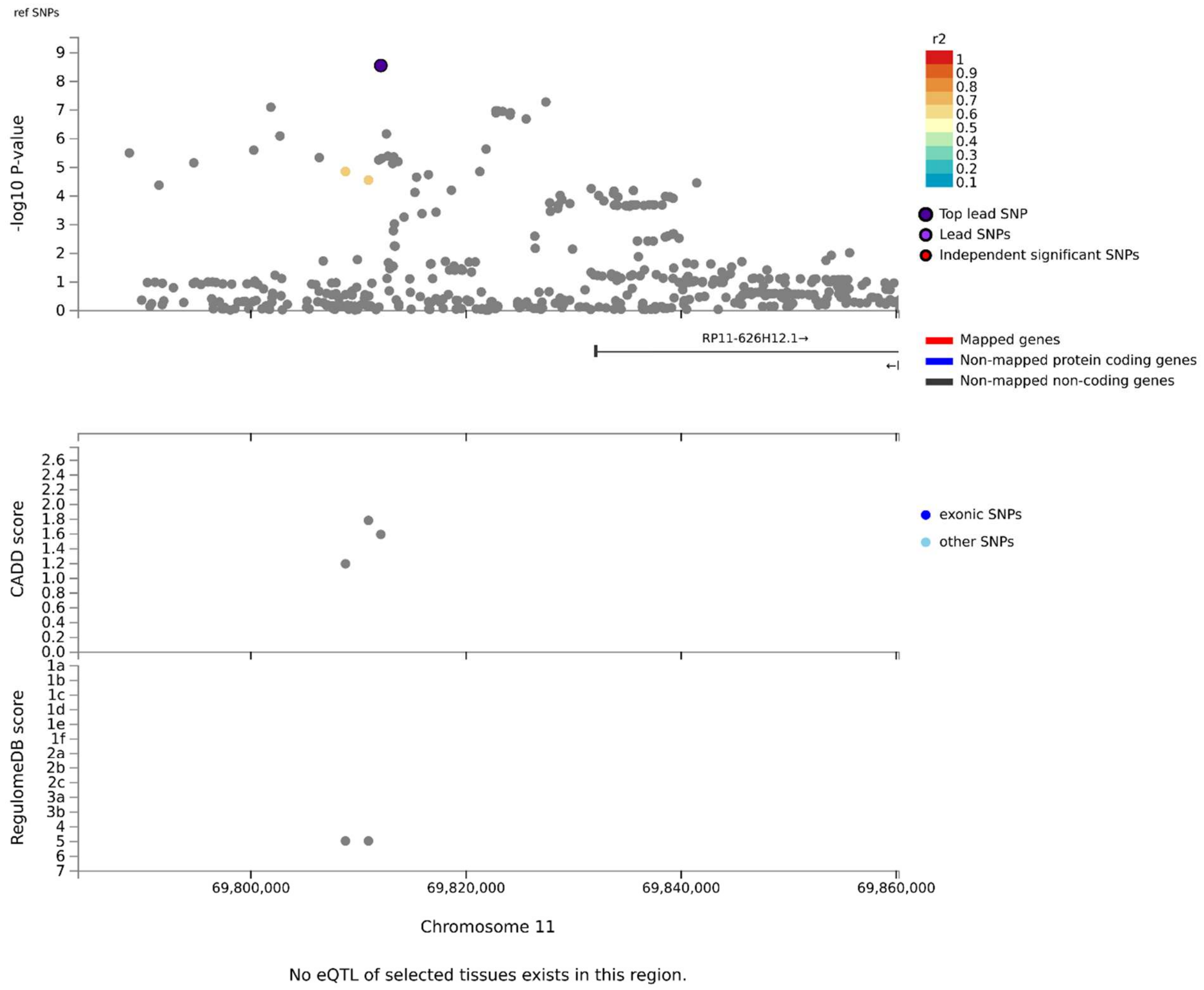


**Supplementary Figure 6.** LocusZoom plot of the locus on chromosome 11 near *RP11-626H12.1* pseudogene with lead SNP rs3929357. The x-axis indicates position on the genome. Top panel indicates association significance with the broad brain language network (-log10 P-values), with the shading indicating linkage disequilibrium. Below nearby genes are shown, then CADD scores (indication of deleteriousness), regulation scores (through RegulomeDB) and associations with gene expression through expressive quantitative trait loci (eQTLs) for the different genes in the neighborhood.

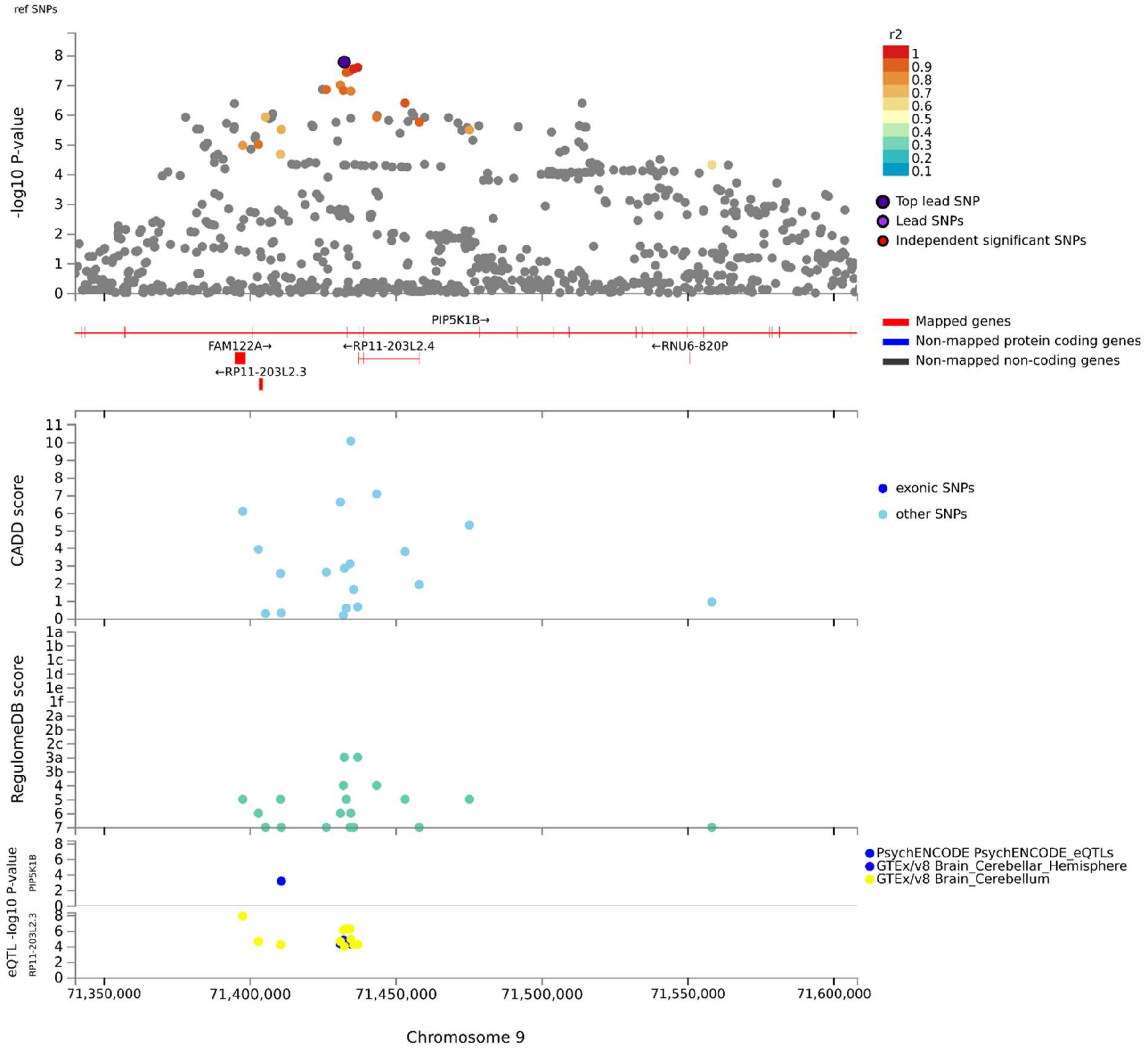


**Supplementary Figure 7.** LocusZoom plot of the locus on chromosome 9 in the *PIP5K1B* gene with lead SNP rs2039425. The x-axis indicates position on the genome. Top panel indicates association significance for the broad brain language network (-log10 P-values), with the shading indicating linkage disequilibrium. Below nearby genes are shown, then CADD scores (indication of deleteriousness), regulation scores (through RegulomeDB) and associations with gene expression through expressive quantitative trait loci (eQTLs) for the different genes in the neighborhood. The lead SNP is close to eQTLs for *PIP5K1B* in the brain.

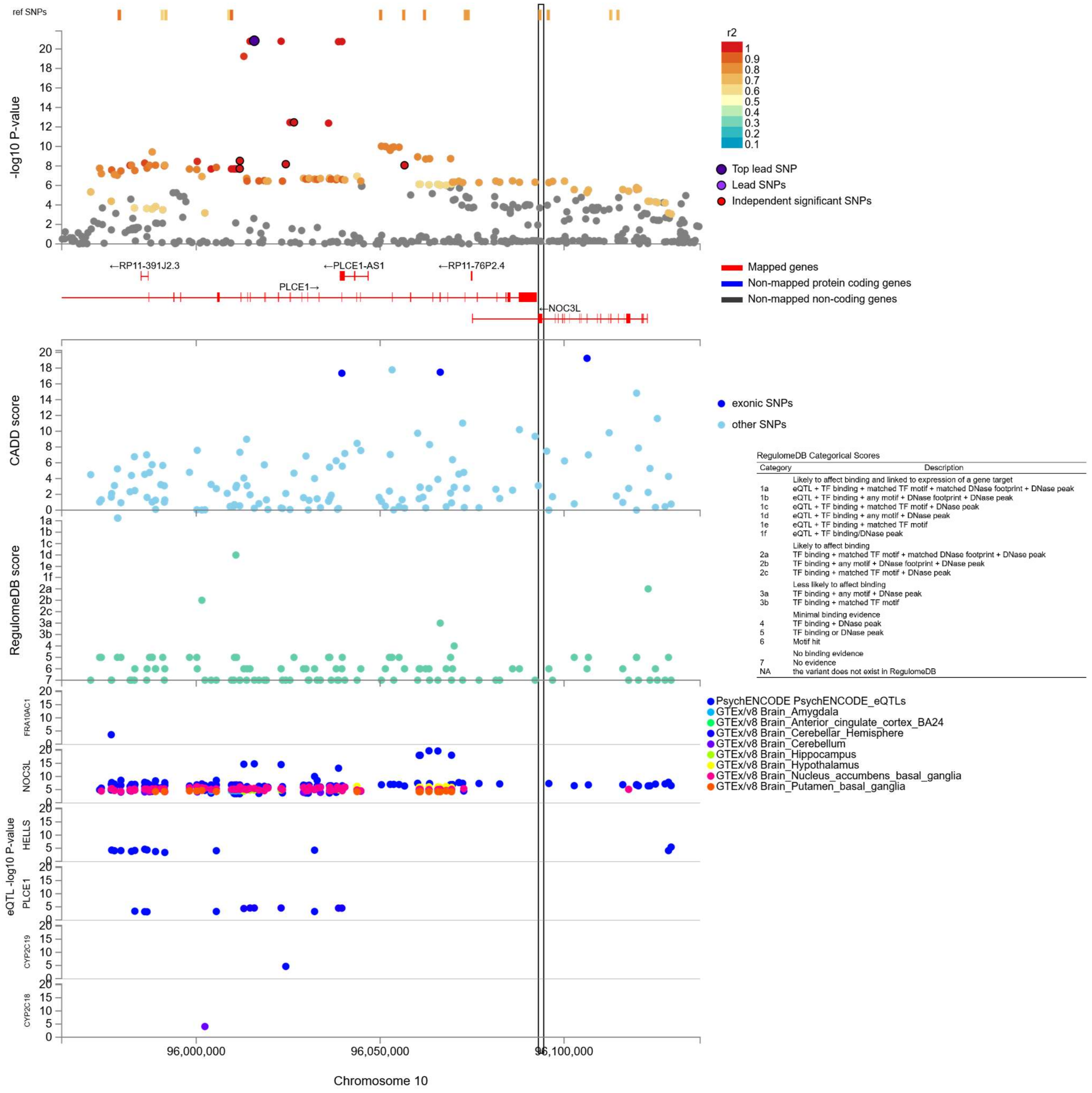


**Supplementary Figure 8.** LocusZoom plot of the locus on chromosome 10 in the *PLCE1* gene with lead SNP rs3891783. The x-axis indicates position on the genome. Top panel indicates association significance with the broad brain language network (-log10 P-values), with the shading indicating linkage disequilibrium. Below nearby genes are shown, then CADD scores (indication of deleteriousness), regulation scores (through RegulomeDB) and associations with gene expression through expressive quantitative trait loci (eQTLs) for the different genes in the neighborhood. The lead SNP is close to eQTLs for *NOC3L* and *PLCE1*.

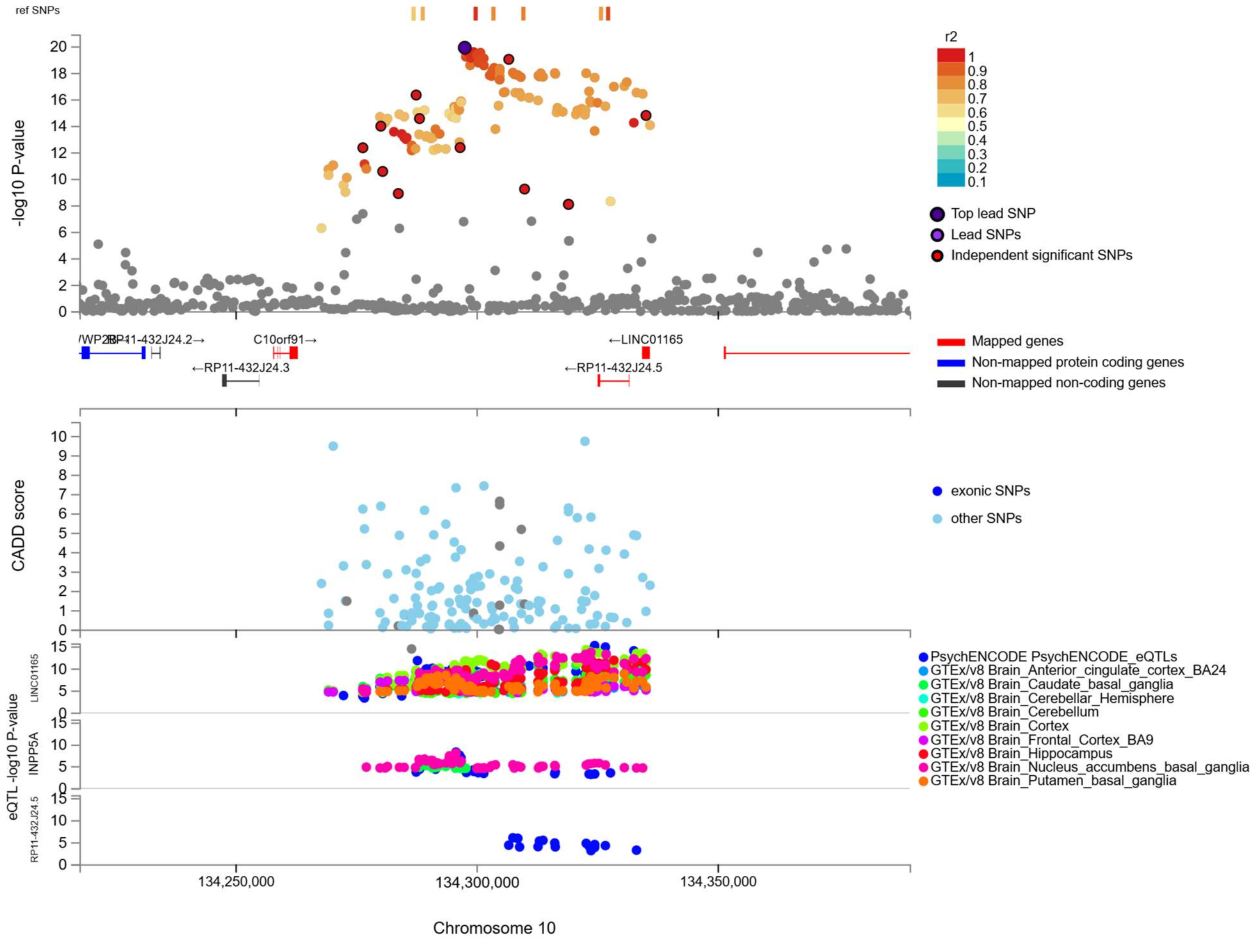

**Supplementary Figure 9.** LocusZoom plot of the locus on chromosome 10 near the *LINC01165* gene with lead SNP rs7084359. The x-axis indicates position on the genome. Top panel indicates association significance with the broad brain language network (-log10 P-values), with the shading indicating linkage disequilibrium. Below nearby genes are shown, then CADD scores (indication of deleteriousness), regulation scores (through RegulomeDB) and associations with gene expression through expressive quantitative trait loci (eQTLs) for the different genes in the neighborhood.

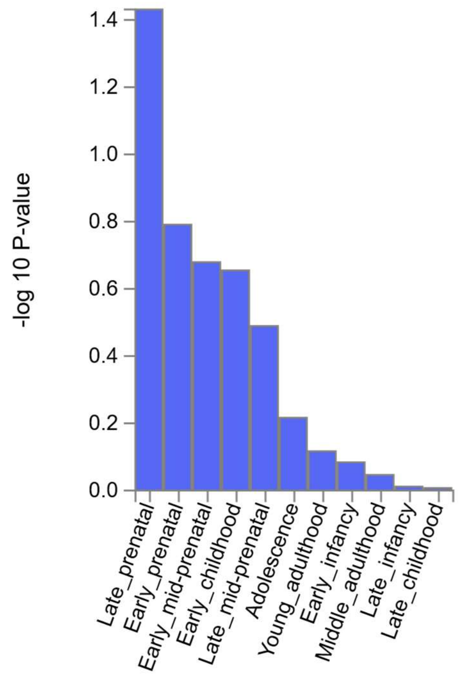


**Supplementary Figure 10.** Genes associated with the broad brain language network component do not show significant expression preferences during a particular developmental stage according to MAGMA analysis of the Brainspan gene expression database.

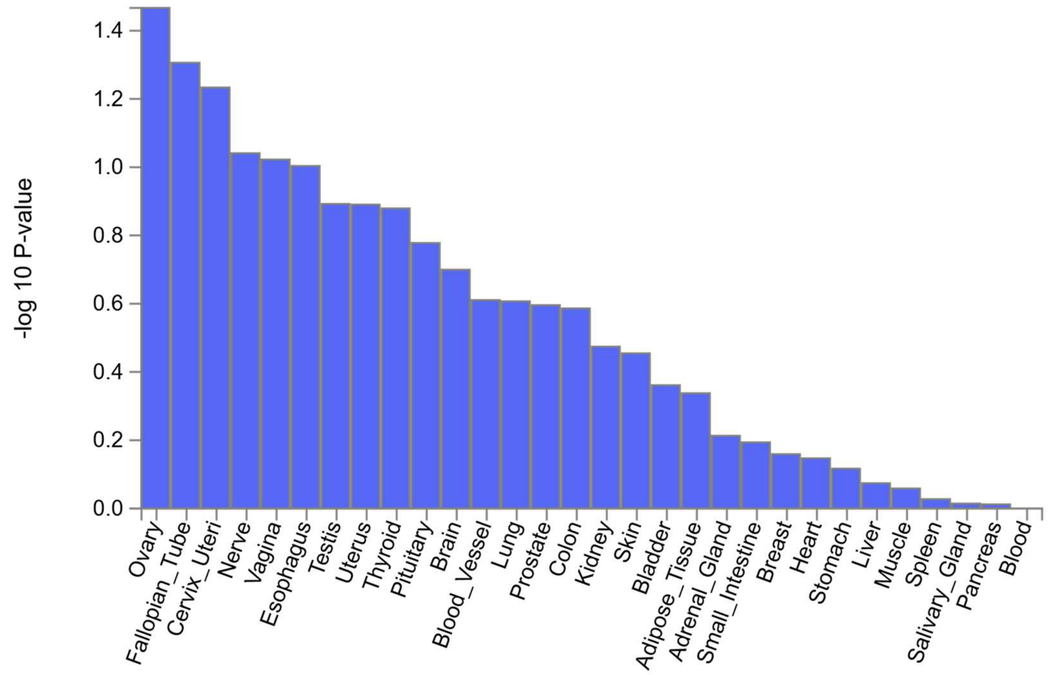


**Supplementary Figure 11.** Genes associated with the broad brain language network component do not show significant preferential expression in a certain tissue type according to MAGMA analysis of the GTEx tissue type database.

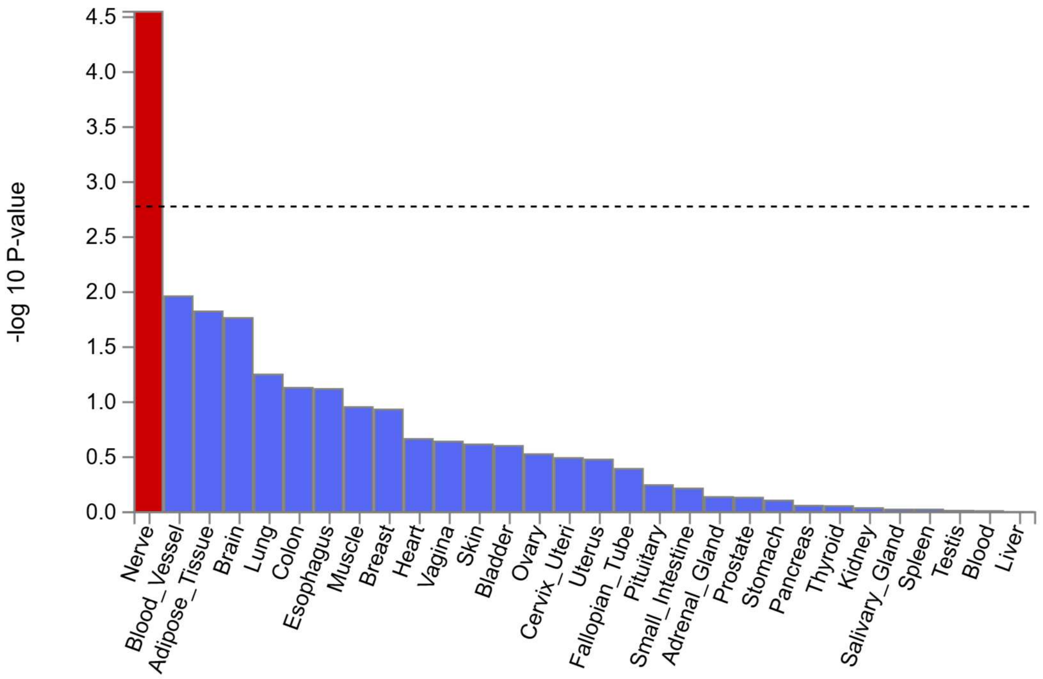


**Supplementary Figure 12.** Genes associated with narrow brain language network component show significant preferential expression in nerve tissue according to MAGMA analysis of the GTEx tissue type database. Dashed line indicates Bonferroni-correction.

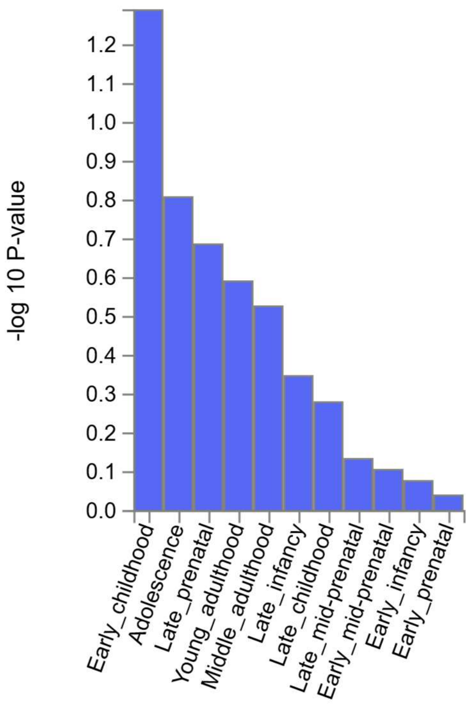


**Supplementary Figure 13.** Genes associated with narrow brain language network component do not show significant expression preferences during a particular developmental stage according to MAGMA analysis of the Brainspan gene expression database.

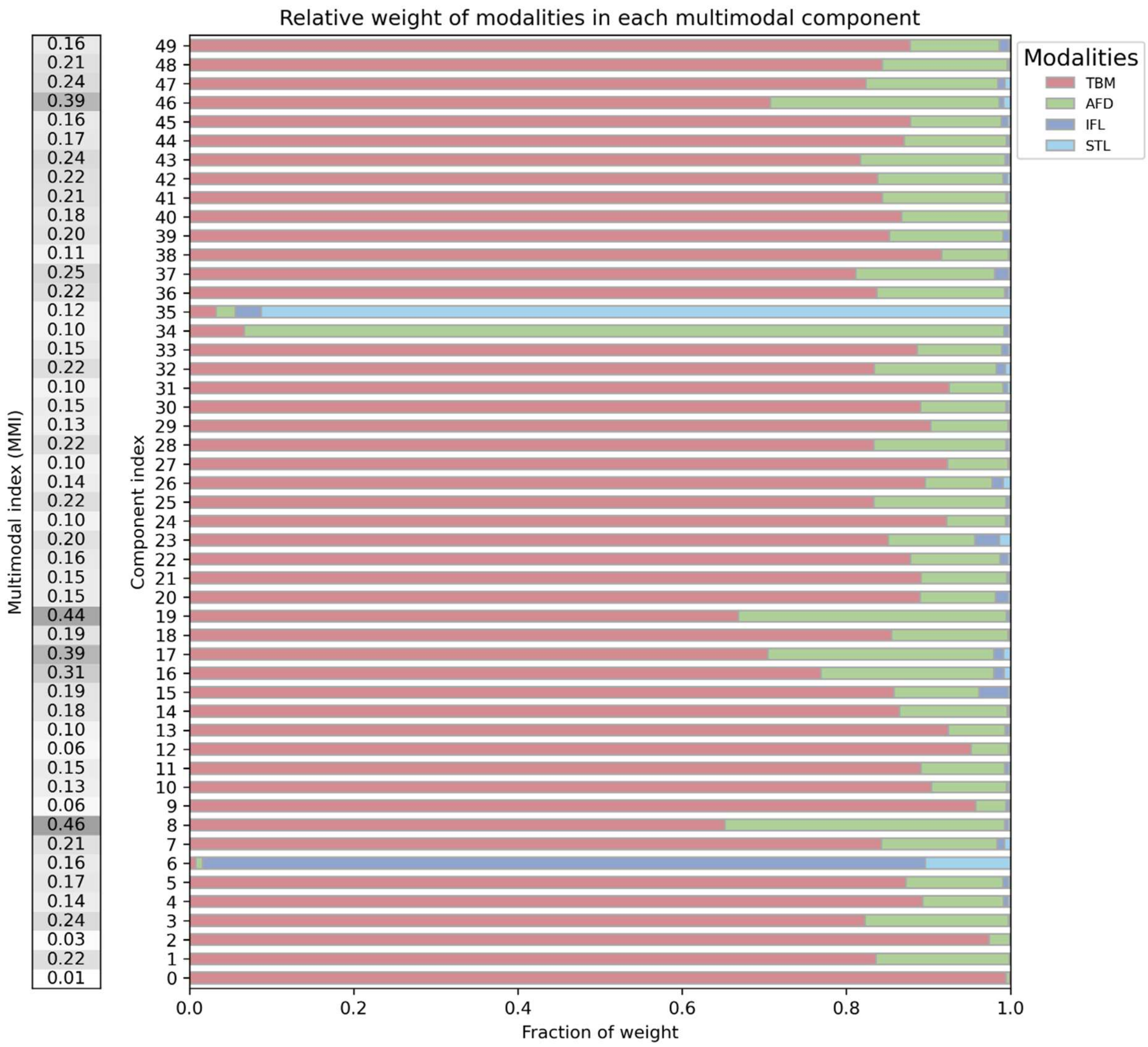


**Supplementary Figure 14.** Fraction weights per modality for all components in the 50 component decomposition. On the left the multimodal index (MMI) is shown, indicative of multimodality. Components 6 (IFL) and 35 (STL) were used in the validation analysis as resting state dominated network components.

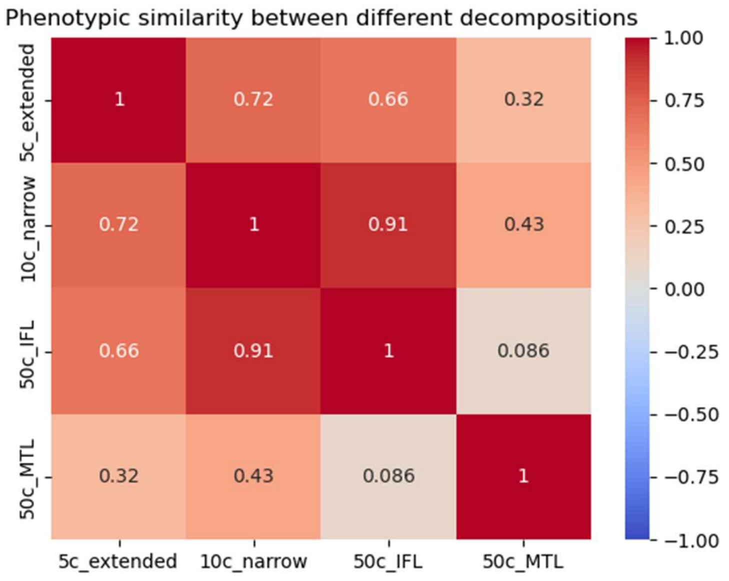


**Supplementary Figure 15.** Phenotypic similarity, uncorrected correlations of subject courses, between different decompositions.

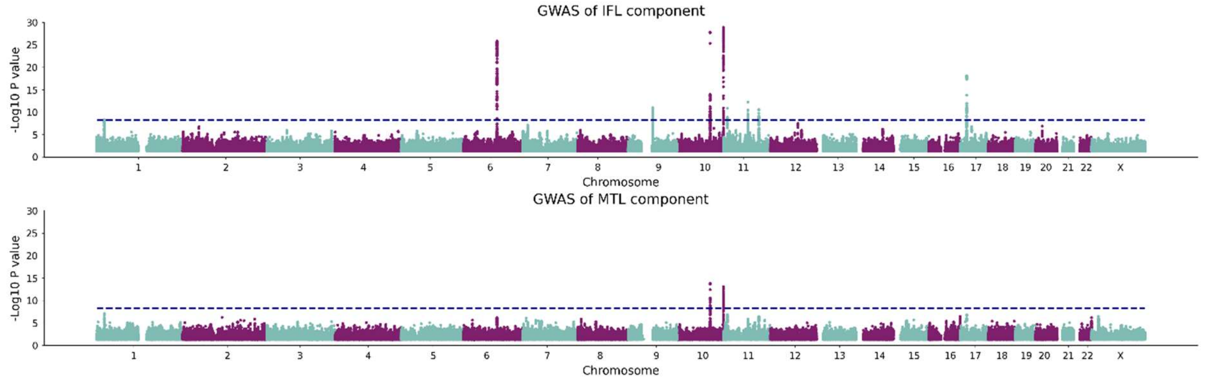


**Supplementary Figure 16.** Manhattan plots for GWAS of components 6 and 35 of the 50 component decomposition, which show largely similar results to the main analyses.
